# Integrin-driven Axon Regeneration in the Spinal Cord Activates a Distinctive CNS Regeneration Program

**DOI:** 10.1101/2021.12.07.471602

**Authors:** Menghon Cheah, Yuyan Cheng, Veselina Petrova, Anda Cimpean, Pavla Jendelova, Vivek Swarup, Clifford J. Woolf, Daniel H. Geschwind, James W. Fawcett

**Affiliations:** John van Geest Centre for Brain Repair, Department of Clinical Neurosciences, University of Cambridge, Cambridge CB2 0PY, UK; Program in Neurogenetics, Department of Neurology, and Department of Human Genetics, David Geffen School of Medicine, University of California, Los Angeles, CA 90095, USA.; Department of Neurobiology, Harvard Medical School; F.M. Kirby Neurobiology Center, Boston Children’s Hospital, Boston, MA 02115, USA; Centre for Reconstructive Neuroscience, Institute of Experimental Medicine CAS, Prague, Czech Republic; Department of Neurobiology and Behavior, University of California, Irvine, CA 92697, USA

## Abstract

The peripheral branch of sensory dorsal root ganglion (DRG) neurons regenerates readily after injury unlike their central branch in the spinal cord. However extensive regeneration and reconnection of sensory axons in the spinal cord can be driven by the expression of α9 integrin and its activator kindlin- 1(α9k1), which enable axons to interact with tenascin-C. To elucidate the mechanisms and downstream pathways affected by activated integrin expression and central regeneration, we conducted transcriptomic analyses of DRG sensory neurons transduced with α9k1, and controls, with and without axotomy of the central branch. Expression of α9k1 without the central axotomy led to upregulation of a known PNS regeneration program, including many genes associated with peripheral nerve regeneration. Coupling α9k1 treatment with dorsal root axotomy led to extensive central axonal regeneration and caused expression of a distinctive CNS regeneration program, including genes associated with ubiquitination, autophagy, endoplasmic reticulum, trafficking, and signalling. Pharmacological inhibition of these processes blocked the regeneration of axons from DRGs and human iPS-derived sensory neurons, validating their causal contributions. This CNS regeneration-associated program showed little correlation with either embryonic development or PNS regeneration programs. Potential transcriptional drivers of this CNS program coupled to regeneration include Mef2a, Runx3, E2f4, Tfeb, Yy1. Signalling from integrins primes sensory neurons for regeneration, but their axon growth in the CNS is associated with a distinctive program that differs from that involved in PNS regeneration.

## INTRODUCTION

Regeneration of axons in the damaged central nervous system (CNS) is a currently unfulfilled requirement for the repair of damage to the nervous system. For successful regeneration, centrally projecting neurons need to be modified to enhance their intrinsic regenerative ability, and their properties need to be matched to the environment through which their axons must grow. How this interplay between extrinsic signals and intrinsic neuronal states is represented in genome-wide transcriptional programs is not well understood, and such an analysis requires a model in which there is extensive regeneration of axons in the CNS.

When the peripheral axon branch of sensory neurons is cut, the axons start to regenerate and a set of genetic and epigenetic changes occur in the cell body leading to the transcriptional upregulation of regeneration-associated genes (RAGs) (Chandran et al., 2016; Curcio and Bradke, 2018; Palmisano et al., 2019; Renthal et al., 2020; Rozenbaum et al., 2018). Expression of this peripheral nerve (PNS) regeneration programme after a conditioning peripheral nerve lesion can enable some sensory axon regeneration in the spinal cord (Chandran et al. 2016)(Neumann and Woolf, 1999). However, cutting the central branch in the dorsal root or spinal cord does not initiate the RAG programme.

Recently an integrin-based strategy has enabled long-distance central sensory neuron regeneration in the spinal cord with functional recovery (Andrews et al., 2009; Cheah et al., 2016; Nieuwenhuis et al., 2018). These approaches are based on the knowledge that axons regenerate through the extracellular matrix (ECM), for which the main receptors are integrins. In the uninjured/developing and damaged CNS, the main integrin ligand in the ECM is tenascin-C (Dobbertin et al., 2010; Rauch, 2004). Yet, α9β1 integrin, the primary tenascin-C receptor is downregulated in neurons after development and not re-expressed after injury. Expression of α9β1 integrin by itself is not sufficient to enable regeneration because integrins are inactivated by Nogo-A and chondroitin sulphate proteoglycans (CSPGs) (Andrews et al., 2009). However, the addition of an inside-out integrin activator, kindlin-1, equips sensory neurons with the ability for extensive regeneration in the spinal cord, overcoming the CSPG- induced deactivation of integrins (Cheah et al., 2016). Thus, α9-kindlin-1 transduced neurons can regenerate their axons for long distances along the dorsal column of the spinal cord, making appropriate connections in the dorsal horn and enabling recovery of sensation and locomotion. However, the molecular programs that underly this remarkable change in internal growth state due to extrinsic signaling are not well understood.

The present study was designed to study the mRNA profile of sensory neurons whose axons were regenerating in the spinal cord driven by α9 integrin and kindlin-1. We profiled the changes in mRNA expression in regenerating (dorsal root axotomized) and non-regenerating (non-axotomized) sensory neurons expressing α9 integrin and kindlin-1 compared with axotomized and non-axotomized GFP controls, neither of which regenerated. This data was subsequently compared with recent profiling studies of peripheral sensory regeneration (Chandran et al., 2016; Renthal et al., 2020; Tedeschi et al., 2016). Four experimental groups were created through injection of either AAV5-fGFP or AAV5-α9-V5 + AAV5-kindlin1-GFP into four cervical (left C5-C8) dorsal root ganglia (DRGs) with and without dorsal root crush. The groups were: i) α9-kindlin-1-GFP with dorsal root crush to enable axon regeneration in the spinal cord (**α9k1-crush**), ii) α9-kindlin-1-GFP with no axotomy and therefore no regeneration in order to examine the effects of activated integrin expression without regeneration (**α9k1-naïve**), iii) GFP with axotomy to control for the effects of AAV injection and axotomy without regeneration (**GFP-crush**), and iv) GFP with no axotomy to control for the effects of AAV injection (**GFP-naïve**). The DRGs were removed after 8 weeks, at which time many axons are actively regenerating in the spinal cord, and GFP-expressing neurons were dissociated and selected by fluorescence-activated cell sorting (FACS), then mRNA purified and profiled by sequencing.

The expression profile of neurons expressing α9-kindlin-1-GFP without axotomy, and therefore not regenerating their axons, showed a remarkable number of expression changes, including upregulation of almost all the mRNAs previously identified as RAGs. This profile represents the effects of signalling from activated integrins in the absence of injury. Comparing this group with the CNS regeneration group that were transduced with α9-kindlin-1-GFP with axotomy, we observed a CNS regeneration programme of mRNAs that was exclusively associated with regeneration in the spinal cord. The groups of mRNAs upregulated in these regenerating neurons were related to ubiquitination, autophagy, endoplasmic reticulum (ER) casein kinases, transcriptional regulators, transport/trafficking molecules, signalling molecules. Overall, the study reveals a distinct CNS regeneration programme of mRNAs expressed during integrin-driven regeneration of axons in the CNS.

## RESULTS

### Expression of integrin and kindlin drives sensory regeneration anatomically

In previous work we showed that expression of α9 integrin and kindlin-1 enables extensive sensory axon regeneration in the spinal cord (Cheah et al., 2016). Here we repeated this experiment, confirming the result. The left C5-C8 dorsal root ganglia (DRGs) were injected with either AAV5-fGFP or AAV5-α9-V5 + AAV5-kindlin1-GFP. In half of the animals the left C5-C8 dorsal roots were crushed. Animals were killed after 8 weeks and sensory projections in the spinal cord were examined. Axons from the DRGs were traced using GFP alone or kindlin-1-GFP. In the spinal cord of α9k1-crush animals, many regenerated axons were seen in the dorsal columns with terminals in the dorsal horn (**Fig. 1A- C**). This replicates our previous regeneration experiment (Cheah et al., 2016). In the GFP-crush animals, there was no regeneration into the spinal cord (**Fig. 1D**). In addition, the spinal cords in the α9k1-naïve group were examined to determine whether expression of α9+kindlin-1 caused abnormal sprouting of axons or terminals. This analysis showed that there was no abnormal sprouting of axons in the dorsal columns, and the projections in the dorsal and ventral horns showed no signs of aberrant sprouting (**Fig. 1E-F**). Expression of α9 integrin and kindlin-1 was therefore sufficient to enable extensive regeneration of crushed central sensory axons into the spinal cord, but it did not cause abnormal sprouting of uncut axons. A schematic summary of the design and results of the *in vivo* regeneration study is shown in **Fig. 1G**.

**Figure 1.**
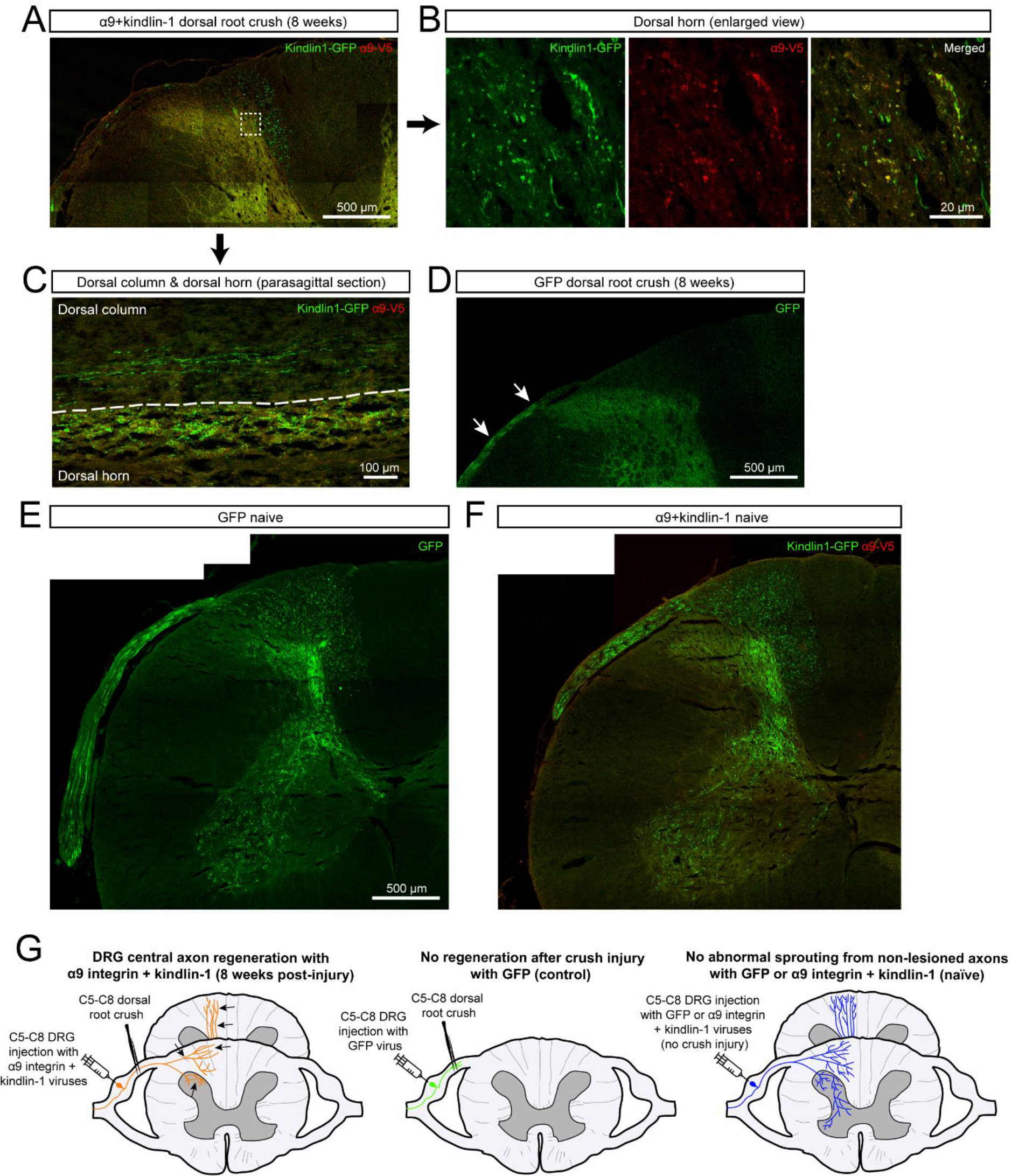
Axonal regeneration and sprouting in the spinal cord. **A** and **B** show regenerated axons in transverse section in the dorsal column and dorsal horn of an α9k1-crush animal. **B** is the magnified square outlined in **A**. **C** shows regenerated axons and their terminals in the dorsal column and dorsal horn in a parasagittal section. For GFP-crush animal in **D**, GFP-labelled axons (*white arrows*) are seen in the dorsal root but none have regenerated into the spinal cord. **E** and **F** show the sensory projections of unlesioned animals, GFP-naive and α9k1-naïve, respectively. There was no abnormal sprouting of axons that contain activated integrin. **G** shows a schematic summary of the design and results of the *in vivo* regeneration study, with regenerating axons (*black arrows*) in the α9k1-crush group.

### Transcriptional changes

DRGs were harvested 8 weeks after virus injection and dorsal root crush, timed to coincide with ongoing axon regeneration. DRGs were dissociated into single cells, then GFP-expressing neurons were selected by FACS, gating for single dissociated cells followed by GFP-expressing neurons (**Fig. S1A**). *Post hoc* sampling showed FACS-sorted cells contained both large and small DRG neurons by co-staining with NF200 and CGRP (**Fig. S1B**).

Transcriptomic analysis via RNA-seq was performed using 4-6 biological replicates for each condition (Methods). Comparison of differentially expressed genes between the GFP and α9k1 groups (Methods) revealed that amongst the most upregulated mRNAs were kindlin-1 (FERMT1, log2FC=6.46) and α9 integrin (ITGA9, log2FC=5.69), demonstrating that AAV transduction for expression of α9 integrin and kindlin-1 had been successful and validating the sorting and sequencing methods. The numbers of differentially expressed genes in the experimental groups are shown in **Fig. S2A**, and the quantitation of overlapping genes in **Fig. S2B-C**.

### Analysis of transcriptional changes by clusters and modules

In our analysis, 2,443 genes were upregulated by integrin-kindlin expression, with or without dorsal root crush, compared to GFP controls (False discovery rate (FDR) of *P* < 0.1; **Fig. S2A**). To identify genes associated with CNS axon regeneration (α9k1-crush, the regeneration group), we applied K-means clustering, identifying 4 clusters, which are visualized on a combined expression heatmap (**Fig. 2**) and shown as individual clusters (**Fig. 3**). Cluster 1 contains genes exclusively upregulated in neurons transduced with α9k1 in combination with central axotomy, many of which were regenerating their axons into the spinal cord (CNS regeneration program). Cluster 2 contains genes upregulated by α9k1 expression without dorsal root crush, but not upregulated in the regeneration group (isolated α9k1 program). Cluster 3 contains genes upregulated in both the α9k1-naïve and α9k1-crush groups (mixed α9k1 program), and Cluster 4 contains genes upregulated by dorsal root crush without α9k1 expression, and therefore without any regeneration (injury program).

**Figure 2.**
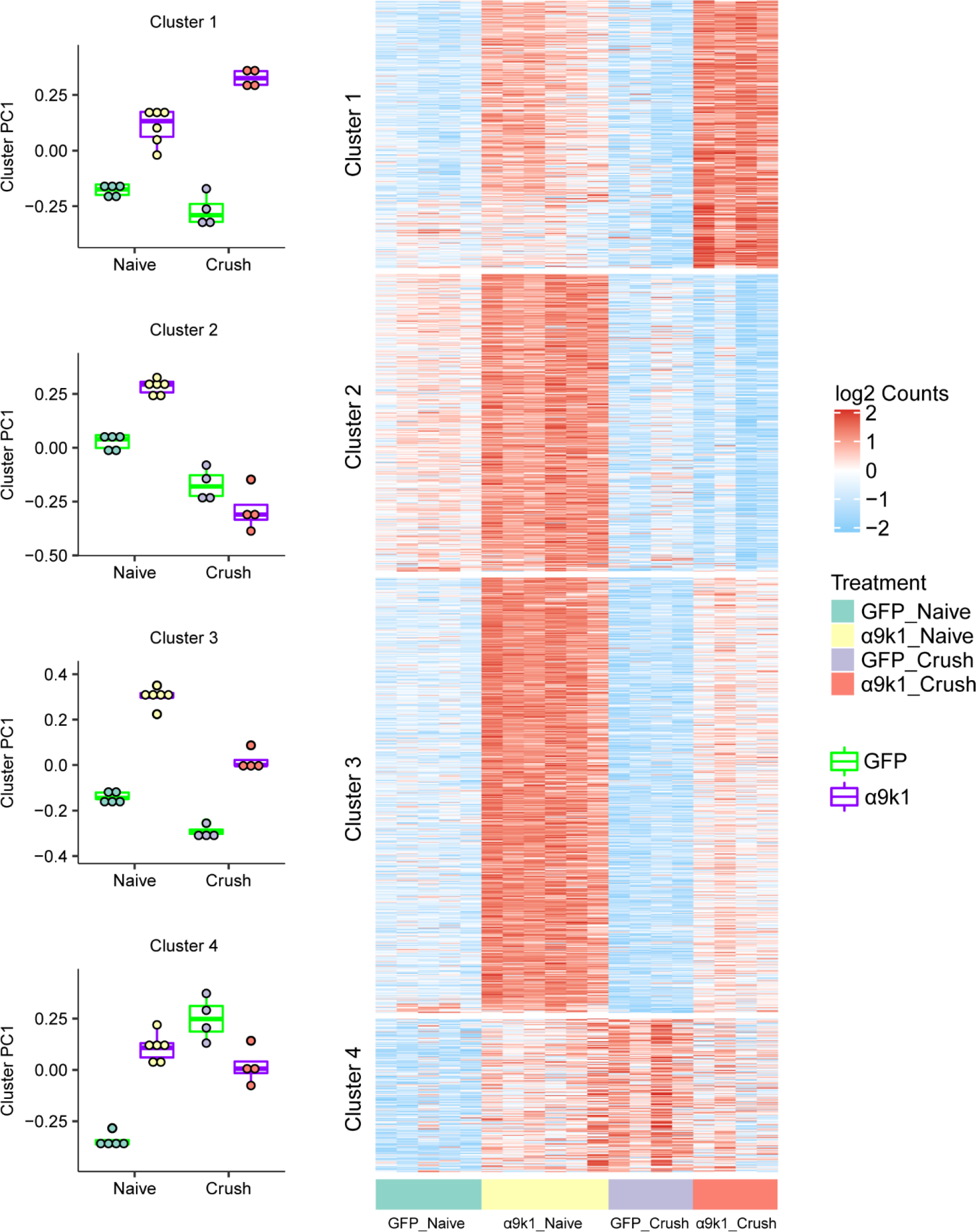
Clustering of genes upregulated by α9k1 expression. K-means clustering was applied to genes differentially upregulated comparing α9k1-naïve to GFP- naïve, or comparing α9k1-crush to GFP-crush (false detection rate, FDR < 0.1). These genes are divided into four clusters, with each associated with α9k1-driven axon regeneration (*Cluster 1*), with α9k1 expression (*Cluster 2*), with α9k1 expression and regeneration (*Cluster 3*) and with crush injury alone (*Cluster 4*). Principal component analysis (PCA) was performed on each gene cluster. The boxplot shows the first PC (PC1), which accounts for the largest variance of gene expression changes against treatment. Expression levels (log2 normalized, row scaled) of individual genes from each cluster were visualized by a heatmap.

**Figure 3.**
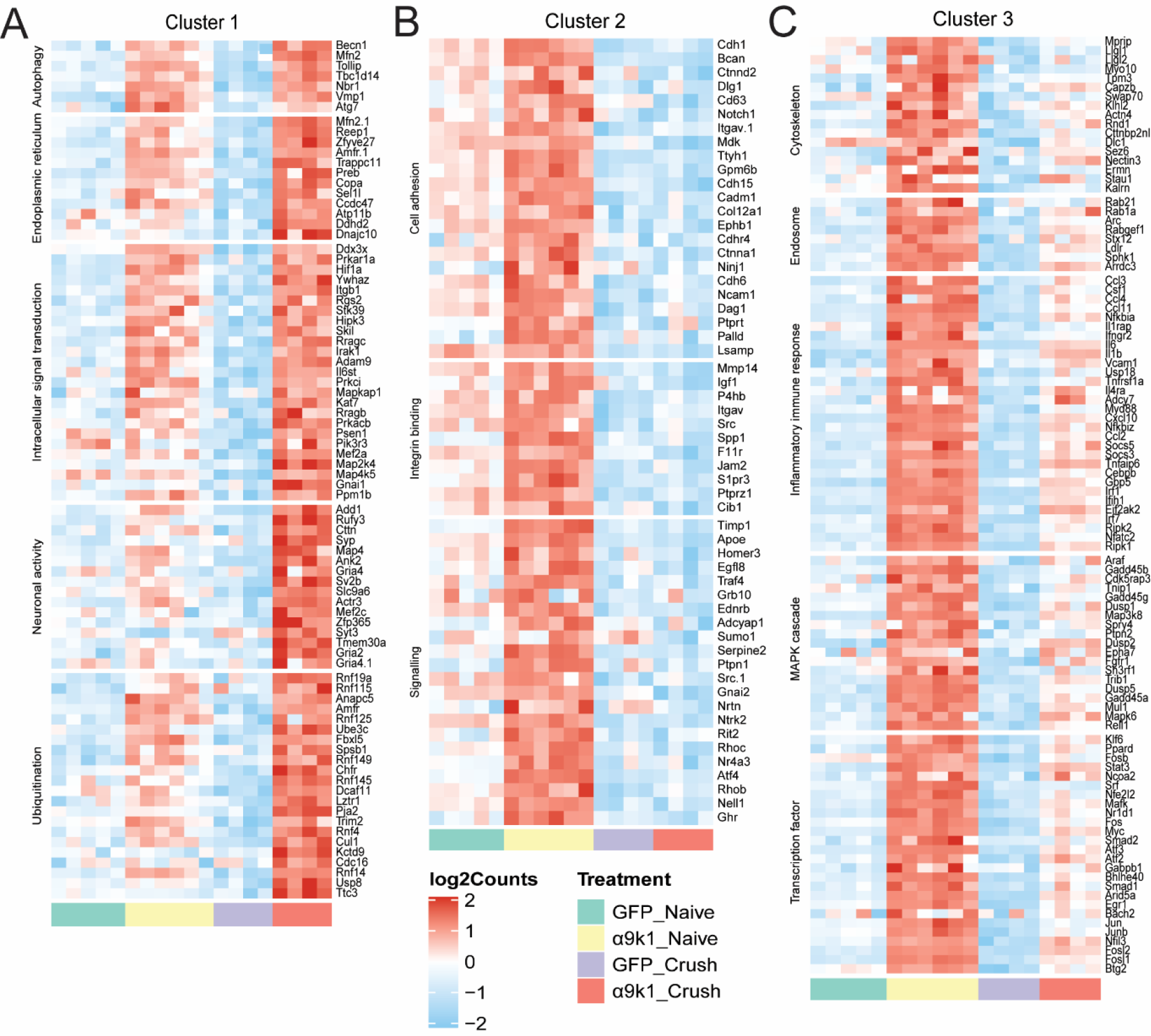
Individual heatmaps with relevant GO terms from Clusters 1,2 and 3. **A** shows Cluster 1 genes upregulated specifically in the α9k1-crush group (*red bar*) with relevant GO terms such as autophagy, endoplasmic reticulum, intracellular signal transduction and ubiquitination. **B** shows Cluster 2 genes upregulated in the α9k1-naïve group (*yellow bar*) with relevant GO terms such as cell adhesion, integrin binding and signalling. **C** shows Cluster 3 genes upregulated in α9k1 groups with or without crush injury (*red and yellow bars*); relevant GO terms include cytoskeleton, endosome, inflammatory response, MAPK cascade and transcription factor.

To further refine our analysis by identifying networks of highly co-expressed genes we performed weighted gene co-expression network analysis (WGCNA) of the entire dataset, identifying (**Fig. 4A**) 11 distinct modules, named by colors as per convention (Langfelder et al., 2008) . Based on the correlation of the first principal component (PC1) of a module (module eigengene) with treatment (Methods; **Fig. 4B**), we found that the Magenta, Black and Green modules were more positively associated with the α9k1-crush (CNS regeneration) group, while the Turquoise and Tan modules were positively associated with the mixed α9k1 program and the Pink module was positively associated with isolated α9k1 program. The relationships of the modules to the various treatment and injury groups (eigengene correlations) are shown in **Fig. 4C** for Magenta, Black, Green, Turquoise, Tan and Pink modules, and in **Fig. S3** for other modules not relevant to our subsequent analyses.

**Figure 4.**
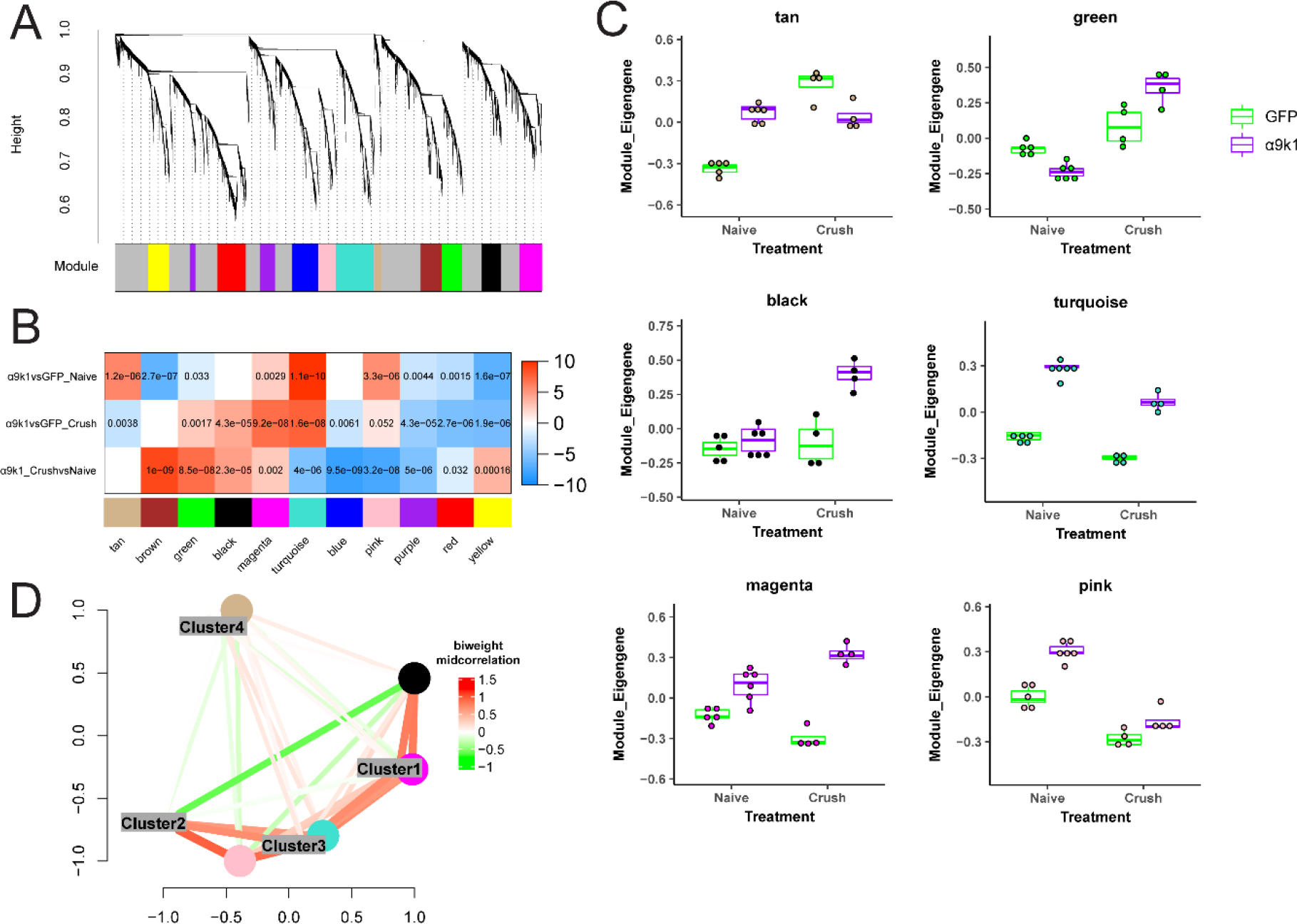
WGCNA of the dataset showing different expression patterns of the modules. **A.** Dendrogram with 11 WGCNA modules. **B.** Correlation between module eigengenes, the first principle component driving the expression changes of a module, with treatments. In the correlation heatmap, colors indicate –sign(correlation coefficient)*(log10 p-value). Red indicates a positive correlation and blue indicates a negative correlation. Numbers shown are Bonferroni-corrected p- values. Green, Magenta and Black modules were more associated with the α9k1-crush (regeneration) group, while Tan, Turquoise and Pink modules were more associated with the α9k1-naïve group. **C.** Trajectory of the module eigengene across treatment and control groups for Tan, Green, Black, Turquoise, Magenta and Pink modules. **D.** Multidimensional scaling plot showing correlations between module eigengenes of the WGCNA modules and DEG clusters (Figure 1). Colors indicate bi- weighted mid-correlation (R) values. This analysis demonstrates similarities between Cluster 1 and Magenta, Cluster 3 and Turquoise.

Comparison of the clusters and modules by similarity network analysis showed that there was a close correlation between the Magenta module and Cluster 1, the Turquoise module and Cluster 3 (**Fig. 4D**). A general Gene Ontology (GO) analysis did not find relevant biological processes associated with the Tan and Green modules. Therefore, the Magenta, Black, Pink and Turquoise modules and equivalent clusters were examined in more detail since they appeared to represent coherent biological processes. The GO of the clusters and the relevant modules were determined (**Fig. 3, S3 and S4**). Genes in GO terms of interest ranked by FDR-corrected P-value were manually curated to select transcripts of potential relevance. Similar GO terms were combined in **Fig. 3A-C and S4** and **Tables S1 and S2**.

A distinctive set of transcripts representing a CNS regeneration program was upregulated only in the α9k1-crush condition (CNS regeneration group), seen in Cluster 1 and the Magenta and Black modules (**Fig. 3A, Fig. 4B-C**). The most significant GO terms included signalling, protein ubiquitination, autophagy, endomembrane system organization, transport/trafficking/localization, vesicle organization, and cytoskeletal protein binding (**Fig. 5 and S4, Table S1**). A different set of transcripts was associated with expression of α9k1 with or without dorsal root crush (mixed and isolated α9k1 programs). These genes appear in Clusters 2 and 3 and in the Pink and Turquoise modules (**Fig. 3, 4B- C, Table S2**) and represent immune processes, cytokine production, transcriptional regulation, biological adhesion, signaling, and ubiquitin-protein transferase activity. Gene networks of these genes and their curated terms are shown in **Fig. 5A-B.** To visualise interactions between the selected genes, network interaction diagrams were made for the hub genes (kME > 0.85) enriched in Pink/Turquoise (**Fig. 5A**) module GO terms and for the Black/Magenta gene networks (**Fig. 5B**). These networks reveal a rich set of interactions that couple the biological processes related to α9k1 signaling in distinct patterns depending on their relationship to central sensory axon regeneration in the spinal cord.

**Figure 5.**
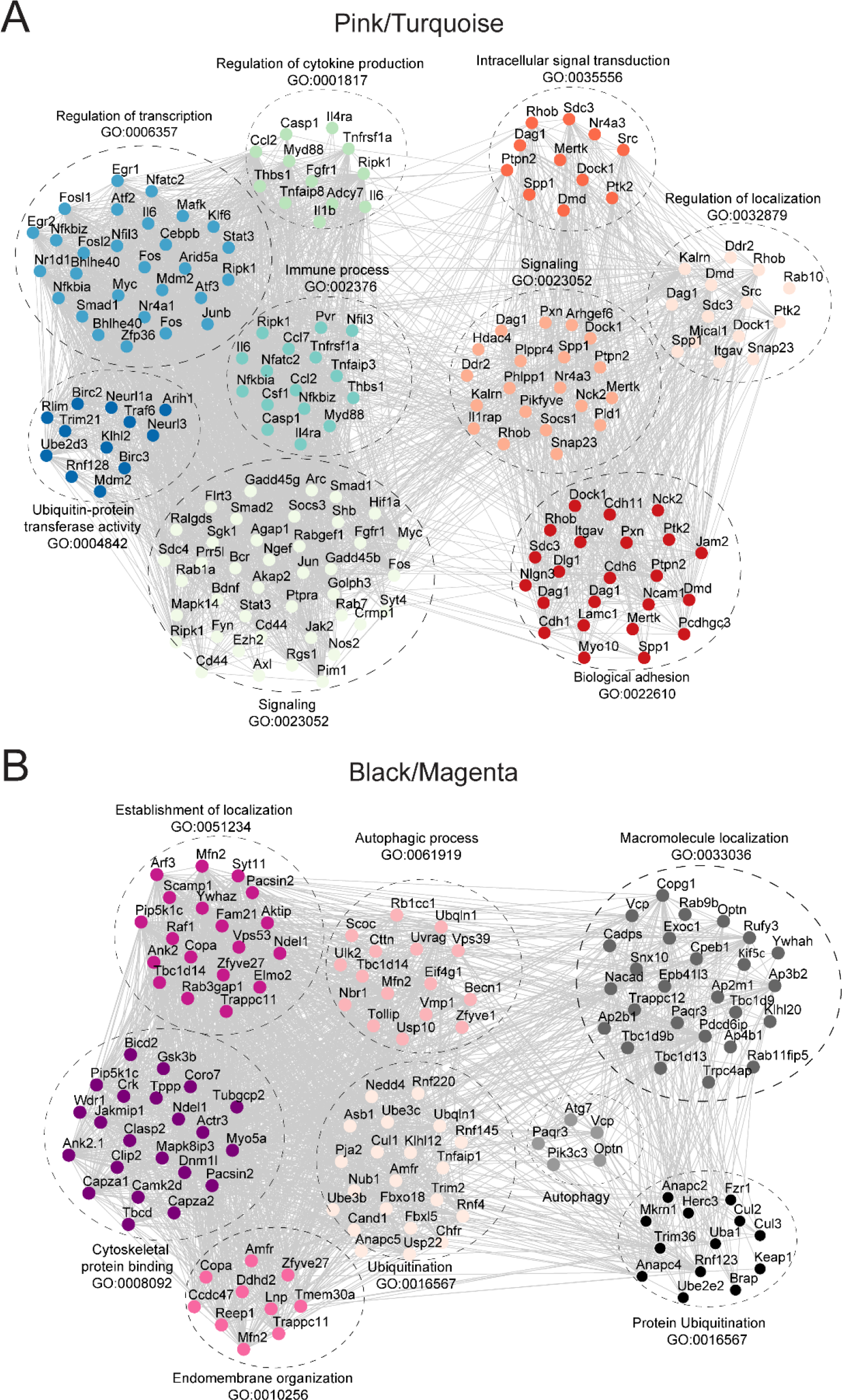
Gene co-expression networks of Pink/Turquoise and Black/Magenta modules. GO analysis was performed for these modules, and co-expression networks were plotted for top- ranked related GO terms. **A.** For the Pink/Turquoise modules, these interaction diagrams show very extensive interactions between the inflammation, transcription, signalling, ubiquitin and localization molecules that were regulated by expression of α9 integrin and kindlin-1, with or without dorsal root crush. **B.** The Gene Ontology analysis of the α9-k1-regeneration-related Magenta/Black modules revealed that genes related to ubiquitinylation, autophagy, and endomembrane organization (with many endoplasmic reticulum-related molecules) had a high level interaction.

### Expression of integrin and kindlin upregulates the PNS regeneration programme

Analysis of the expression level clustering and co-expression network module analyses were performed to examine correspondence between transcripts in Clusters 2-3 and Pink and Turquoise modules that were upregulated by α9k1 expression (**Fig. 3B-C, S4, Table S2**). Cluster 2 includes mRNAs related to cell adhesion, integrin binding, and signalling (**Fig. 3B**). Cluster 3 GO terms of interest include actin skeleton, early endosomes, inflammatory immune response, MAPK cascade, and transcription factors (**Fig. 3C, Table S2**).

The transcription factor term in Cluster 3 is of particular interest since it contains many of the traditional regeneration-associated molecules (RAGs), including the recognized transcription factor RAGs KLF4, KLF6, Jun, Myc, ATF3, EGR1, EGR2, Smad1, STAT3, Fos, Fosl1, Fosl2, CEBP. A larger list of regeneration-related mRNAs was identified in Chandran et al. (Chandran et al., 2016) (*Chandran RAGs list*). Many of these appear in Cluster 3 and Pink and Turquoise modules in the transcription factor, inflammation and signalling GO terms (Transcripts that appear in our Cluster 3, Pink and Turquoise, and also in the Chandran et al. RAGs module are highlighted in green in **Table S2**).

Given this overlap, we next asked whether the integrin-driven changes resemble those that occur during PNS regeneration. Over-representation analysis revealed a strong correlation between the pink and turquoise modules and genes upregulated during PNS regeneration in a previous study (Chandran et al., 2016) (**Fig. 6A**). A correlation analysis using rank-rank hypergeometric overlap (RRHO) heatmaps compared the logFCs of the experimental groups vs. GFP controls, and a previous study of mRNA changes after sciatic nerve crush, comparing DRG neurons FACS-purified 1 and 5 days after axotomy with controls (GSE188776). There was a strong correlation between changes in the α9k1-naïve group and changes after PNS axotomy (**Fig. 6B**). Dorsal root crush injury alone, comparing the GFP-naïve with the GFP-lesion group did not upregulate any of these factors. (T**able S2**). These findings indicate that signalling downstream of activated integrins is sufficient to activate a regeneration program. This regeneration program leads to a pattern of mRNA expression that closely resembles that caused by peripheral nerve axotomy and includes most of the major recognized RAGs.

**Figure 6.**
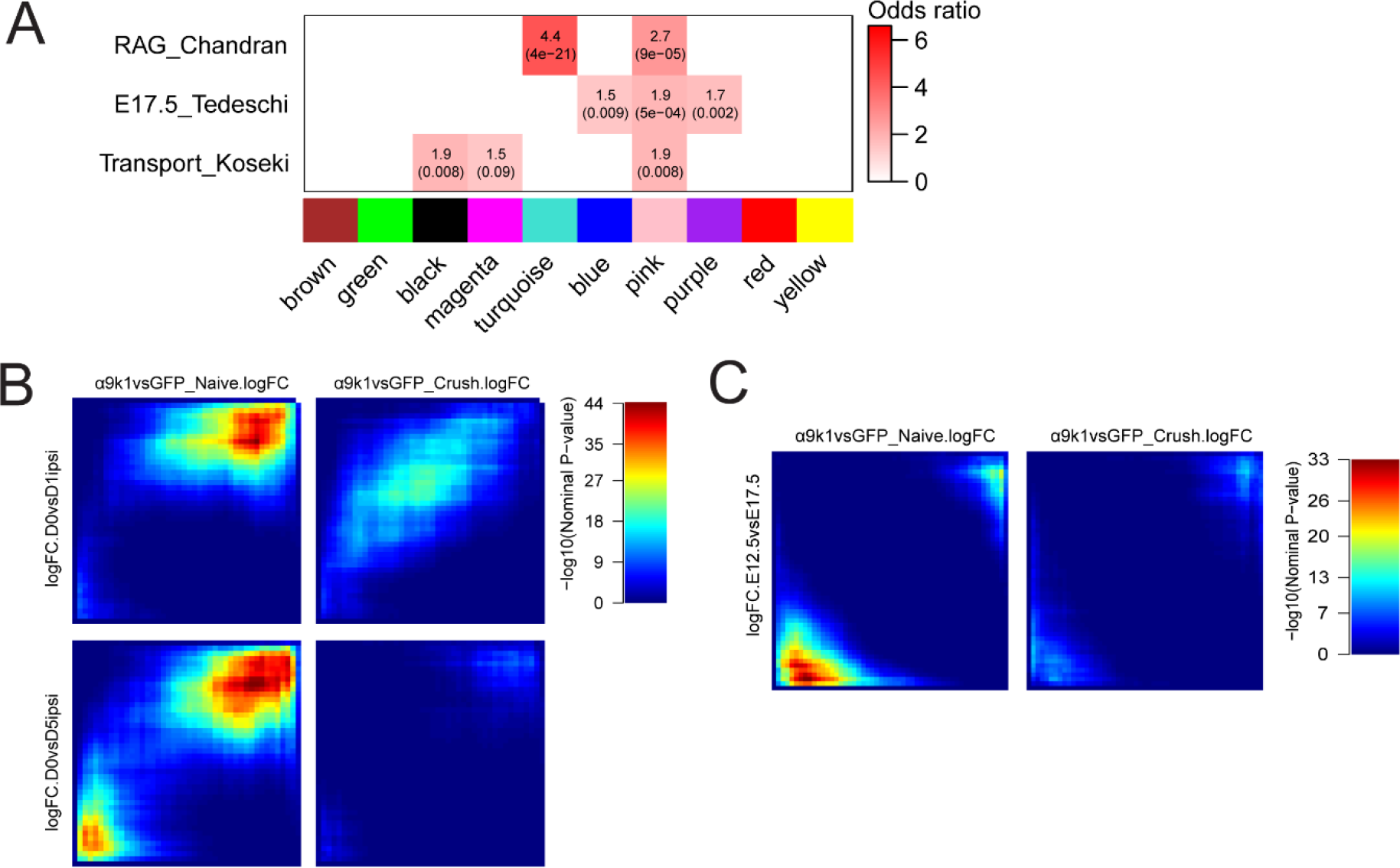
Correlations with PNS regeneration and development. **A.** Gene overlap analysis was performed to determine whether modules regulated by α9k1 are enriched with known gene signatures previously published. Comparisons are between a gene set activated during peripheral nerve regeneration (RAG_Chandran; Chandran et al., 2016), genes associated with DRG neuronal development (E8_Tedeschi; Tedeschi et al., 2016), or the genes associated with neuronal trafficking (Transport_Koseki; Koseki et al., 2017). In the enrichment heatmap, numbers shown are odds ratio indicating the possibility of enrichment, with hypergeometric p-value in parenthesis. **B.** RRHO maps comparing ranked logFC of genes from the α9k1 data and the logFC of same genes from DRGs with sciatic nerve injury). The α9k1-naive group is correlated with the peripherally injured DRGs, especially the downregulated genes on both time points, 1 and 5 days after axotomy (*top right corner*). Each pixel represents the overlap between genes from two different datasets, color-coded according to the -log10 p-value of a hypergeometric test (step size = 200). On each map, the extent of shared upregulated genes is displayed in the top right corners, whereas shared downregulated genes are displayed in the bottom left corners. **C.** RRHO maps comparing ranked logFC of genes from the a9k1 data and the logFC of same genes from embryonic DRG neurons at E12.5 and E17.5. However, there were no significant correlation between any of the experimental groups, and genes expressed during embryonic axon growth.

### Spinal cord regeneration is associated with upregulation of a CNS regeneration programme

A distinct set of genes was associated only with the α9k1-crush (CNS regeneration) group. We observed that most of the genes described above that were upregulated in the α9k1-naïve group also remained upregulated in the α9k1-crush group (**Fig. 2**). However, we observed another set of genes specifically upregulated only in the α9k1-crush group. These transcripts were found in Cluster 1 (**Fig. 2, 3A**), and in the Magenta and Black modules (**Fig. 4B-C**). Gene ontology analysis of these clusters and modules shows that many of these genes are enriched in biological processes related to autophagy, ubiquitination, endoplasmic reticulum and signalling (**Fig. 3A**, **5B, S4, Table S1**). Transcripts curated for relevance are shown in the network diagram (**Fig. 5B**) and the table of mRNAs associated with GO terms (**Table S1**). Comparison with a previous network of molecules related to axonal transport and trafficking also showed strong correlations, revealing a set of densely networked molecules mostly associated with endosomal transport (**Fig. 6A** and **S5**) (Koseki et al., 2017). Network analysis revealed that these groups of molecules were highly co-expressed (**Fig. 5B**). Interestingly, in contrast to the Pink and Turquoise modules, very few of the genes in these regeneration-related cluster and modules were recognized RAGs (highlighted in **Table S2**), apart from the kinase Camk2d, vesicle-related Pdcd6ip, autophagy-related Tbc1d14 and ubiquitin-related Usp4 and Lnx1 which appear in the Chandran et al. RAGs list. In addition, two casein kinases, CSNK1D and CSNK1A1, were upregulated in the α9k1-crush condition. Association of casein kinases with axon regeneration has been reported previously (Ayad et al., 2016).

### Comparison with gene expression during development

While regeneration shares mechanisms with embryonic development, the molecules involved are often different and spinal cord repair does not closely recapitulate development (Harty et al., 2003; Nacu and Tanaka, 2011; Seifert et al., 2019). A recent profiling study of peripheral nerve regeneration has found only minor correlation between the changes after peripheral axotomy and embryonic development (Renthal et al., 2020). However, another study of expression changes in cortical neurons regenerating their axons into spinal embryonic grafts found a partial recapitulation of the embryonic gene expression pattern (Poplawski et al., 2020).

We compared overlap between our modules with embryonic expression in sensory neurons in the database of Tedeschi et al. (Tedeschi et al., 2016) using Fisher’s exact test (**Methods**). The Pink module showed modest overlap with E17.5 DRG expression in the initial eigengene correlation module analysis (**Fig. 6A**). We also compared the logFCs of genes in the experimental groups with gene expression in embryonic DRGs at E12.5 and E17.5 (Tedeschi et al., 2016) using the RRHO analysis (**Fig. 6C**). There was no significant correlation between any of the experimental groups, and genes expressed during embryonic axon growth (logFC E12.5 vs E17.5). Of the ubiquitin-related and autophagy genes upregulated in our regeneration group, there were no or minor changes between adult vs embryonic in the Tedeschi database. Thus, we observed only modest recapitulation of the embryonic expression pattern following α9k1 expression and regeneration after crush injury.

### Transcriptional Regulation of the PNS and CNS regeneration programmes

To identify potential TF activators and repressors that might regulate the PNS and CNS regeneration programs, we performed a TF enrichment analysis in the promotors of the genes in key regeneration associated modules (**Methods**). We aimed to identify TFs that are both present and have regulatory targets in the modules representing the PNS and CNS programs. For the PNS regeneration program there are many potential regulators. Many TFs in the RAGs module from Chandran et al. are regulated by integrin expression and have binding sites in the Pink and Turqoise modules, including Jun, JunB, Egr1, Rela, Crem, Atf2, Atf3, Fos, Cebpd, Runx2, Stat3 and Bhlhe40. However, fewer transcription factors were found in the Black and Magenta modules representing the CNS regeneration program. As potential regulators of ubiquitination, autophagy, ER, localization and cytoskeletal binding genes that appear in the modules associated with the CNS regeneration programme we identifed Mef2a, Runx3, E2f4, Tfeb, Yy1 (**Fig. 7A**). TF enrichment analysis predicts many connections between these TFs and the CNS regeneration program targets (**Fig. 7A**).

**Figure 7.**
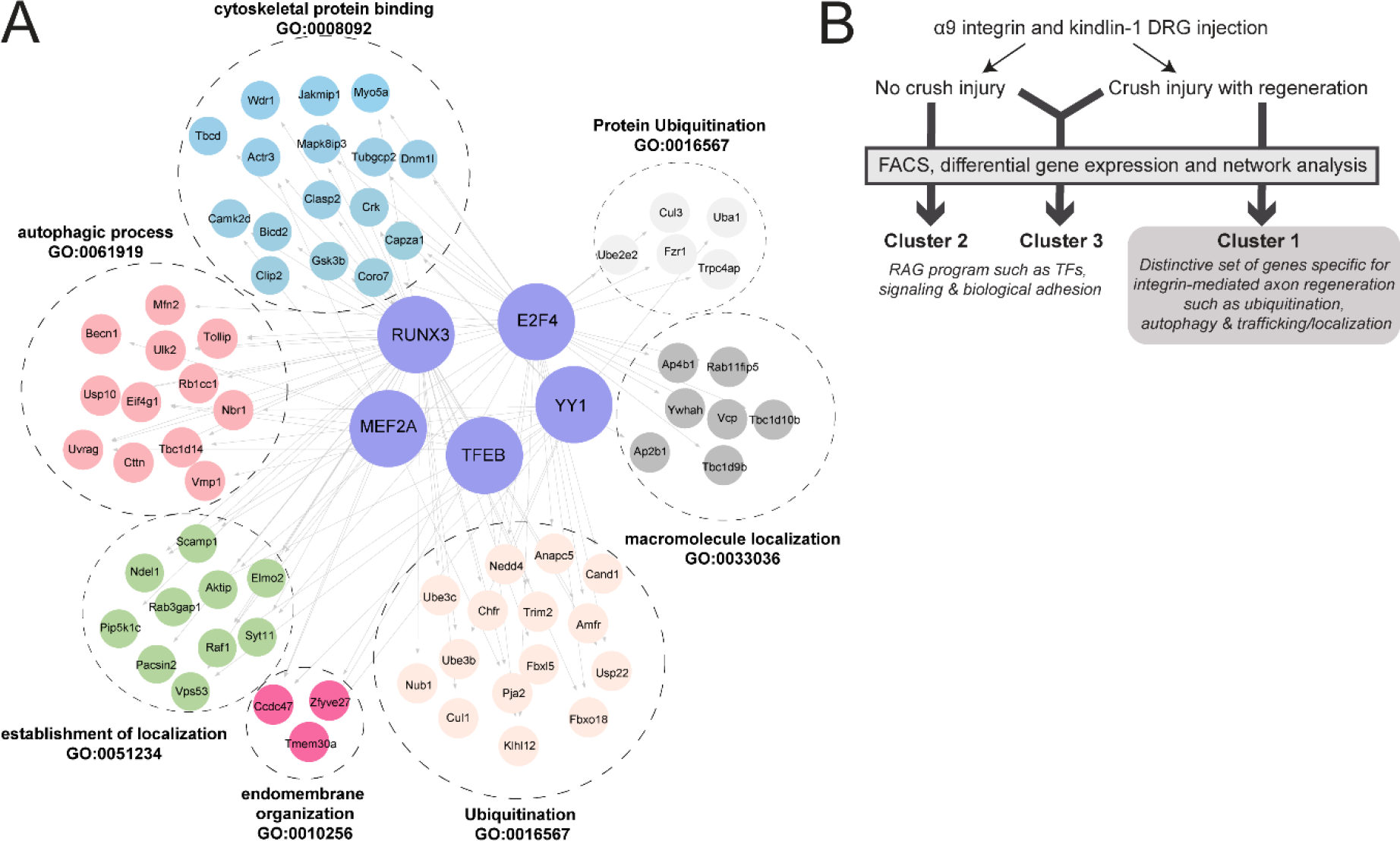
Transcriptional Control of the CNS regeneration program. **A.** TF enrichment analysis identified five potential transcriptional controllers of the CNS regeneration program: Runx3, E2F4, Yy1, Tfeb and Mef2A. These transcription controllers could regulate ubiquitination, autophagy and localization processes. **B.** Schematic diagram summarizing the main findings of this study where we identified a distinctive set of CNS regeneration program (Cluster 1) which could play a role in integrin-mediated axon regeneration.

### Central sensory nerve regeneration is associated with upregulation of ubiquitin, autophagy and ER-related mRNAs

A novel finding of this study was the upregulation of many ubiquitin-related molecules in the α9k1- crush (CNS regeneration) group, seen in Cluster 1 and Magenta/Black modules (**Fig. 3A, 5B, S4, Table S1**). Many of these molecules are related to ubiquitination via SCF complexes (SKP, Cullin, F-Box), specifically E3 ubiquitin ligases that attach ubiquitin to molecules via the K48 and K11 lysine residues and associated adaptor/recognition molecules (marked yellow in **Table S1**) (Oh et al., 2018). Based on the analysis, 15 of the upregulated ligases (marked grey in **Table S1**) ubiquitinate via the K63 position.

This form of ubiquitination is commonly associated with modulation of function of enzymes and trafficking of molecules and receptors (Oh et al., 2018).

UbiNet (Nguyen et al., 2016) was utilized to search for ubiquitination networks and hubs amongst mRNAs expressed in the Magenta/Black/Turquoise modules as these modules were highly upregulated in the α9k1-crush (CNS regeneration) group (**Methods**). This revealed four main ubiquitination networks (**Fig. 8 and S6**). The cullin family and Btrc are the nodes of a network (**Fig. 8A**) associated with ubiquitination of molecules targeted to the proteasome, including the SCF complex together with SKP1 and F-Box proteins, and interacting E3 ligases. Molecules believed to interact with cullins are outlined in yellow in **Tables S1 and S2**. A second smaller network has Nedd4 and Nedd4L as hubs (**Fig. 8B**). These ubiquitin ligases have already been associated with axon growth through interactions between PTEN and mTORC1 (Christie et al., 2012; Drinjakovic et al., 2010; Hsia et al., 2014). A further network (**Fig. S6A**) centres around the hub Traf6, an E3 ligase implicated in promoting tumorigenesis and invasion through activation of AKT signalling (Feng et al., 2014; Yang et al., 2009) and in control of NF-kappa-B. A fourth network (**Fig. S6B**) is associated with oxidative stress and neuroprotection. It has Ube2e2 and Keap1 as hubs: Ube2e2 is an E2 ubiquitin conjugating enzyme, while Keap1 is a substrate-specific adaptor that senses oxidative stress acting through the cullin-3 complex (Kobayashi et al., 2004).

**Figure 8.**
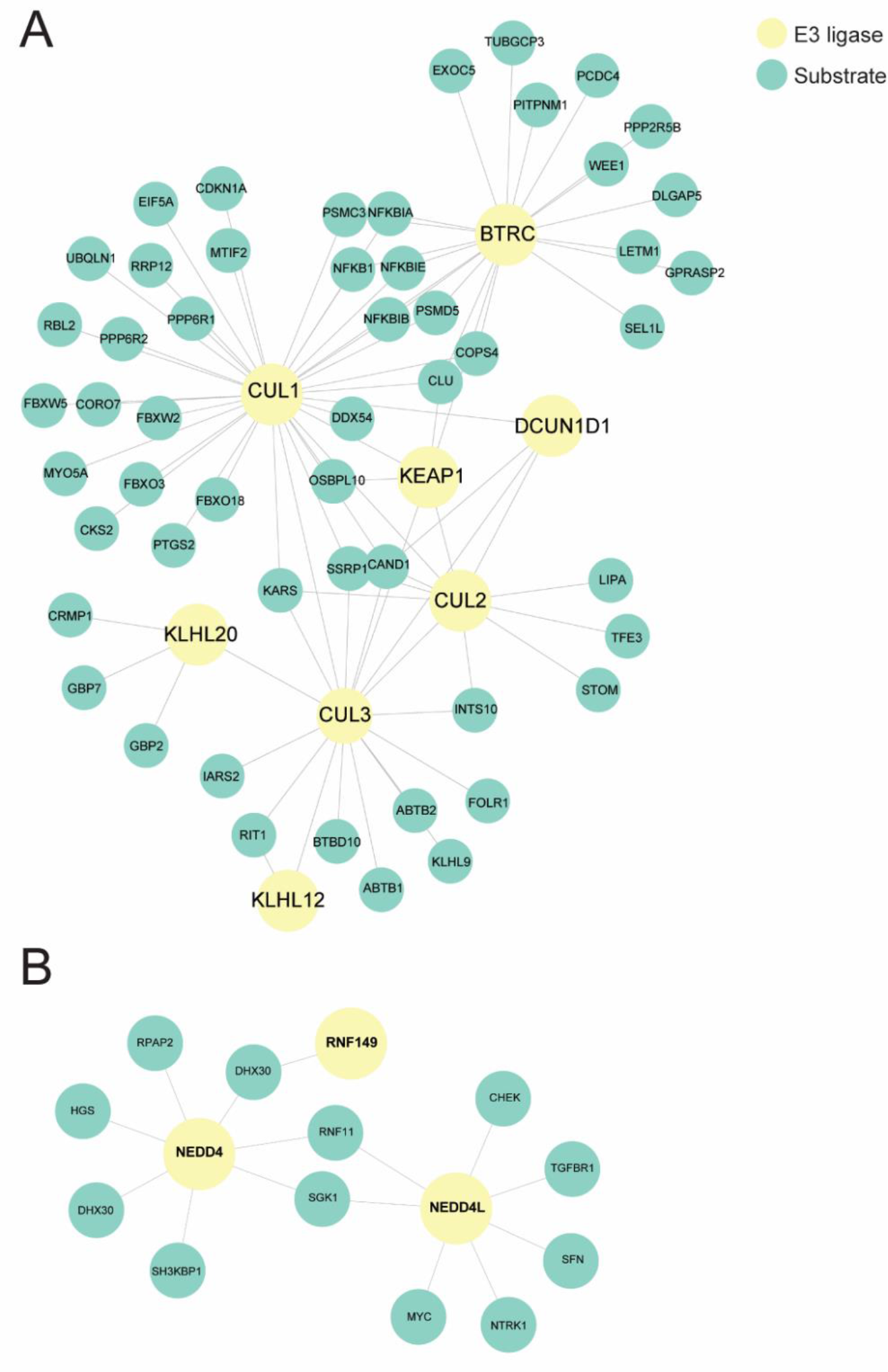
Analysis of ubiquitin-related mRNAs upregulated during sensory regeneration using UbiNet. **A.** An extensive network centers around the cullin family and Btrc, which is associated with ubiquitination of molecules targeted to the proteasome. **B.** A smaller network with Nedd4 and Nedd4L as hubs, which could potentially be associated with regulation of axon growth via interactions with PTEN and mTORC1.

Autophagy was a significant term in Cluster 1 and the Magenta module. Only three of the Atg genes which code for autophagosome structural moleculles were upregulated. But, there was upregulation of several transcripts coding molecules that play a key role in the initiation and control of autophagy (**Table S1**). Optineurin, NBR1 and FIP200 are autophagy receptors that recognize ubiquitin chains; ULK, Beclin, VMP1, PAqR3 and Pik3c3 form a complex that generates local PI(3)P recognized by dfcp1 to initiate autophagosome formation; LC3 and Gabarap are part of the autophagosome membrane; VMP1 is ER-linked and participates in autophagosome formation and VCP in autophagosome maturation (Dikic, 2017; Kuijpers et al., 2020; Kulkarni et al., 2018).

### Inhibition of ubiquitination, autophagy and casein kinases affect in vitro axon regeneration

The pattern of expression after α9k1-crush (CNS regeneration) suggests that ubiquitination and autophagy are involved in central axon regeneration, and two Type-1 casein kinases were also upregulated. To test whether these processes are necessary for adult sensory axon regeneration, we applied specific inhibitors to explants of adult rat DRG explants whose axons were allowed to grow for 5 days before axotomy (**Fig. 9A-B**). We also tested regeneration after axotomy from human iPSC-derived sensory neurons (Singec et al., 2020) that were grown as spot cultures after 21 days of maturation, and whose axons were axotomized with a laser 400-600 µm away from the cell body (**Fig. 9C-E**). In the rat DRG explants, regeneration of cut axons began after 20 mins and the percentage of regenerating axons was measured at 2 hours (**Fig. 9A**); continuing growth of uncut axons was also measured (**Fig. 9B**). Regeneration of the human iPSC-derived sensory neurons was slower, and regeneration was assayed at 6-24hrs (**Fig. 9C-E**). In these assays spontaneous regeneration is already maximal in rat DRG explants (70-80% of axons), and as high as 50-70% in human iPSC-derived sensory neurons, so they are useful for testing potential inhibitors of regeneration. In previous studies on rat DRGs, interventions that block regeneration may have little effect on the continuing growth of uncut axons (Chierzi et al., 2005; Verma et al., 2005).

**Figure 9.**
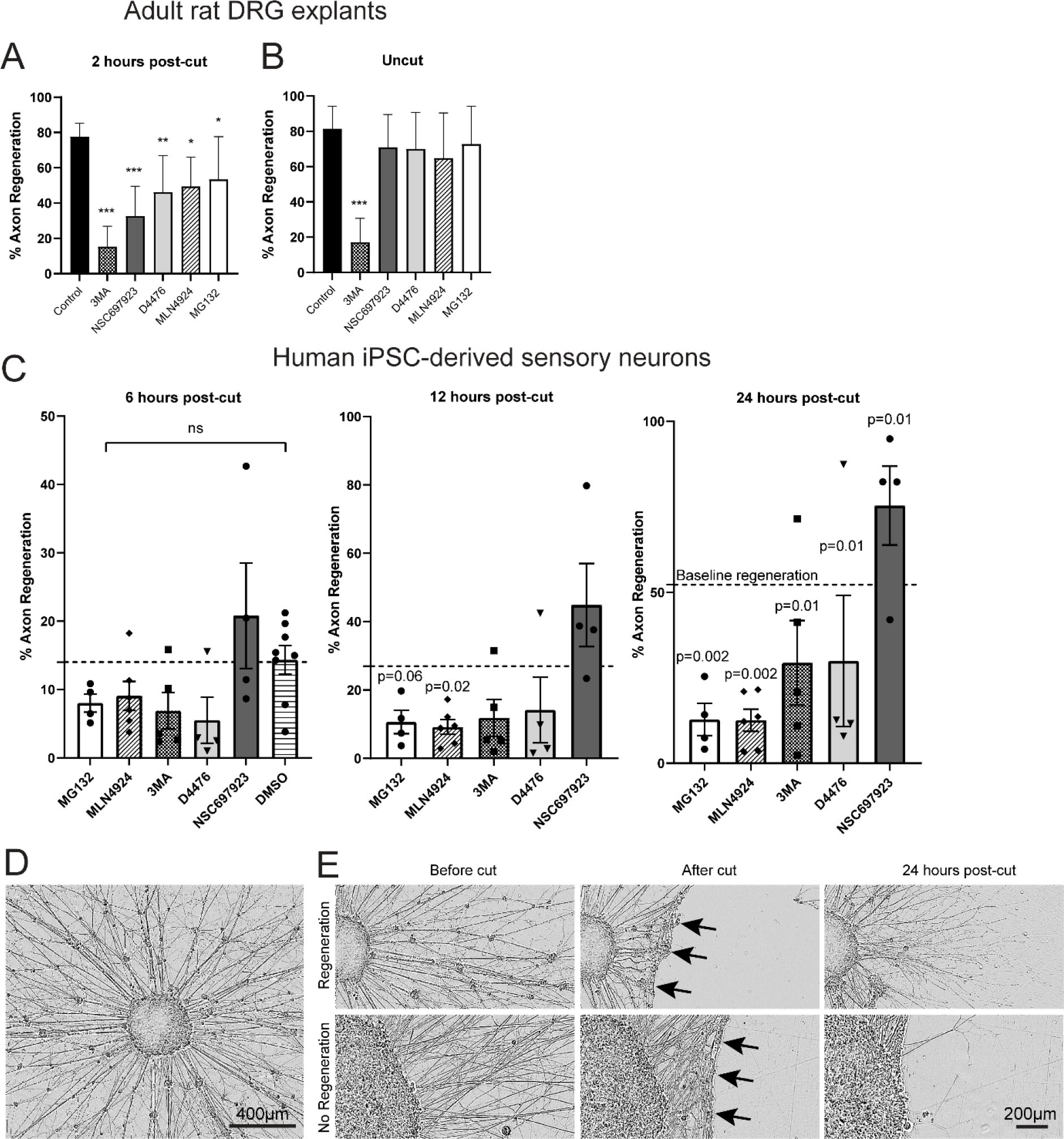
*In vitro* sensory axon regeneration of adult rat DRG explants and human iPSC-derived sensory neurons after axotomy in the presence of inhibitors. The inhibitors used were **MLN492** - NEDD8-mediated ubiquitination inhibitor; **NSC697923** - K63- ubiquitination inhibitor; **MG132** - proteasome inhibitor; **3MA** - Class III PI3K autophagy inhibitor; **D4476** - casein kinase 1 inhibitor **A-B.** For axotomised axons (**A**) in adult rat DRG explants, all the five inhibitors exhibited various degrees but significant inhibition to axon regeneration, with the greatest effect observed by 3MA and the most modest effect by MG132. For uncut axons (**B**), all inhibitors, except 3MA, did not affect continuous axon growth. Analysis was performed using one-way ANOVA with *post hoc* test. A *p*-value of < 0.05 was considered to be statistically significant. \**p*-value < 0.05, \*\**p*-value < 0.01, \*\*\**p*-value < 0.001. **C** shows the percentage of human iPSC-derived sensory axons regenerating at 6, 12 and 24 hours after laser axotomy, and regeneration was inhibited by all the inhibitors except NSC697923. Analysis was performed using Fisher’s exact test. The *p*-values were then analyzed with the “Analyze a stack of *p*- values” function in GraphPad Prism with a Bonferroni-Dunn pairwise comparison. **D** shows an aggregate of human iPSC-derived sensory neurons with halo of axons, after 21 days of maturation. **E** demonstrates the results of laser axotomy of the halo of axons. After axotomy, the axons retracted (*black arrows*) followed by various degrees of regeneration in the presence of inhibitors in the post- 24 hours.

The inhibitors used in these two rat and human in vitro assays were: **MLN492** (inhibits NEDD8 activating enzyme and blocks ubiquitination associated with the SKP,Cullin, F-box complex); **NSC697923** (blocks K63-type ubiquitin ligases by blocking the E2 conjugating enzyme); **MG132** (proteasome inhibitor); **3MA** (inhibits class 3 PI3Ks which locate omegasome [autophagosome precursor] production to the surface of the ER); **D4476** inhibits Casein Kinase 1. In the rat DRG explants, all inhibitors partially blocked regeneration, with the strongest effects from the inhibition of autophagy and E2 conjugating enzyme (**Fig. 9A**). However only inhibition of autophagy affected continuing axon growth without axotomy (**Fig. 9B**). In the human iPSC-derived neurons, inhibitors of autophagy, degradative proteasome, ubiquination and casein kinases all inhibited regeneration, but blocking K63-type ubiquitination did not (**Fig. 9C**).

## DISCUSSION

The mechanisms by which extrinsic signals influence neuronal intrinsic programs that enable regeneration in the CNS are not well understood. Here, we leveraged a model whereby integrin expression is modulated, so as to enable long-distance axon regeneration in the CNS central branches of DRG sensory axons, to begin to elucidate the intrinsic growth programs. We were able to deconvolute expression changes caused by integrin expression alone from those caused by integrin-driven regeneration by comparing two groups with very different outcomes: 1) in the α9k1-naïve group integrin signalling occurred, but the intact sensory axons and terminals in the spinal cord showed no sign of enhanced sprouting; 2) in the α9k1-crush group, there was an extensive axon regeneration along the cord and terminations in the dorsal horn exactly as seen previously (Cheah et al., 2016). We conclude that signalling from activated integrins primes the neurons, putting them in a regeneration-competent state, very similar to that seen in peripheral nerve regeneration after a peripheral axotomy. However when the α9k1-primed neurons are axotomized and regenerate their axons through the CNS environment, activation of a second and previously undiscovered program is required, whose purpose is probably to enable the interaction between the axons, the integrins and their environment.

### Alpha9 integrin-kindlin-1 expression upregulates the PNS regeneration programme

A well characterized program including many transcripts is upregulated as injured peripheral sensory axons regenerate in the PNS (The RAGs program) (Chandran et al., 2016; Rozenbaum et al., 2018; Tedeschi et al., 2016; van Kesteren et al., 2011). Several of these RAGs participate in regeneration (Fagoe et al., 2015; Mehta et al., 2016). A surprising finding was that a substantial fraction of known the PNS regeneration RAG program was induced by α9k1 expression alone (**Table S2**), including most of the RAGs transcription factors. These changes in expression must have been due to signalling from integrins. The signalling would have come not only from the α9 integrin that was virally expressed, but also from the other integrins (of which adult DRG neurons express several) which would be activated by kindlin-1. Integrins signal via focal adhesion kinase and integrin-linked kinase activates the PI3K/Akt, Ras/Erk, Jnk, Rho/Rac pathways and via mechano-transduction through Piezo/Ca^++^, and Hippo/Yap/Taz (Humphries et al., 2019; Hynes, 2002). The unexpected conclusion is that the RAGs programme can be activated by signalling from integrins without any actual axon injury or regeneration.

### Alpha9 integrin-kindlin-1 driven regeneration upregulates a CNS regeneration group of genes

By comparing the α9k1 dorsal root crush and naïve groups, we were able to identify genes that were specifically associated with central axon regeneration in the spinal cord, present as Cluster 1 in the heatmap (**Fig. 2, 3A**) and in the Magenta/Black modules (**Fig. 5B**). In these regeneration-associated cluster and modules, the most significant GO terms were related to transport/trafficking, ubiquitination, autophagy, endoplasmic reticulum, endomembrane organization, signalling and cytoskeletal binding (**Fig. 3, 4, 5, S5 and Table S1**). These processes are presumably involved in bringing molecules to the tips of growing central axons, both enabling the dialog between the growing axons and their environment and supplying the intrinsic growth mechanisms required for growth cone function.

*Transport and trafficking* upregulation in axon regeneration have been recognized for many years, (Grafstein and Murray, 1969; Oblinger et al., 1987). The molecules needed to build growing axons come partly from the cell body via axonal transport, and partly from local translation within axons (Koley et al., 2019; Smith et al., 2020). While transport within CNS neurons is highly selective to direct dendritic postsynaptic molecules and axonal presynaptic molecules correctly, PNS neurons are less selective (Britt et al., 2016; Franssen et al., 2015; Koseki et al., 2017; Maeder et al., 2014). In our CNS regeneration gene group, transcripts upregulated were associated with molecular motors, endocytosis, exocytosis, vesicle sorting, scaffolding, ArfGAPs, Golgi and ER. Two scaffolding molecules, Ywhaz (14.3.3 protein) and Zfyve27 (Protrudin) are of particular interest because both are associated with the promotion of CNS axon regeneration, and Protrudin achieves this by linking trafficking to the ER (Kaplan et al., 2017; Petrova et al., 2020). The transcripts relating to signalling and cytoskeletal binding link to a previous literature on mechanisms of axon regeneration (He and Jin, 2016; Mahar and Cavalli, 2018; Murillo and Sousa, 2018).

*Ubiquitination* molecules of several types were present in the regeneration-associated Cluster 1 (**Fig. 3A**) and Magenta/Black modules (**Fig. 5B**). Many were related to the degradative ubiquitination-proteasome pathway in which E3 ligases associate with the SCF complexes (SKP, Cullin, F-Box), attaching ubiquitin to molecules via the K48 and K11 lysine residues of ubiquitin (Dikic, 2017; Oh et al., 2018; Zinngrebe et al., 2014). Within this group there were cullins, E3 ligases and adaptors. K63 ubiquitination does not usually lead to degradation, but can modulate the function of signalling and transport molecules and marks molecules for trafficking (Oh et al., 2018); there were several E3 ligases of this type. We used UbiNet to search for potential ubiquitylation networks (Nguyen et al., 2016), identifying four networks with hubs emerged (**Fig. 8, S6**). **1)** Cullins are the key component of the SCF box complex, and molecules ubiquitinylated by this mechanism are usually targeted for proteasomal degradation (**Fig. 8A**) (Oh et al., 2018). In combination with the substrate adaptor KLH20, cullin-3 has been implicated in neurotrophin-induced axon growth through degradation of RhoGEF (Lin et al., 2011). **2)** Nedd4 and Nedd4L hubs (**Fig. 8B**). Nedd4 has been implicated in axon growth, although the mechanism is disputed between targeting PTEN for degradation, or as a target of PTEN and mTORC1 (Drinjakovic et al., 2010; Hsia et al., 2014). **3)** E3 ligase Traf6 as hub (**Fig. S6A**). Traf6 controls NF-kappa- B levels, Jun and PI3Kinase-AKT signalling (Hamidi et al., 2017), and also controls autophagy (Nazio et al., 2013). **4)** Keap1 and the E2 ligase Ube2e2 as hubs (**Fig. S6B**), which acts as an oxidative stress sensor (Kobayashi et al., 2004), potentially providing neuroprotection.

*Autophagy* control molecules were also present in the α9k1-crush (regeneration) group. These include ULK1 and Beclin-1, controllers of autophagosome production, and Pik3c3, the kinase involved in omegasome production, a precursor of autophagosomes (Ktistakis, 2020; Maday and Holzbaur, 2016; Winckler et al., 2018). Selective autophagy links the ER, recycling endosomes and ubiquitination, all implicated in axon regeneration (Eva et al., 2012; Petrova et al., 2020), and ubiquitinated molecules are recognized by autophagy recognition molecules such as NBR1, FIP200 and optineurin (present in our Magenta and Black modules, **Fig. 5B**) (Hara et al., 2008; Kenific and Debnath, 2016; Mowers et al., 2017). Autophagy can act as a recycling pathway for integrins and focal adhesions in cell migration (Abbi et al., 2002; Kenific and Debnath, 2016; Kenific et al., 2016; Maday and Holzbaur, 2016; Sharifi et al., 2016; Tuloup-Minguez et al., 2013; Wei et al., 2017). This pattern of ubiquitin molecule expression in the CNS regeneration programme is not seen in PNS regeneration studies (Chandran et al., 2016; Tedeschi et al., 2016), suggesting that this pattern of ubiquitination and autophagy is required specifically for axon growth within the inhibitory CNS environment.

*Endoplasmic reticulum*-related genes were also upregulated. The presence of endoplasmic reticulum in axon growth cones is a requirement for successful regeneration (Petrova et al., 2020; Raiborg et al., 2015; Rao et al., 2016). Endoplasmic reticulum is a source of autophagosomes, generation being controlled by ULKs, Beclin and PIK3C (Dikic, 2017; Stavoe and Holzbaur, 2018), which were upregulated in our regeneration group.

### Validation of involvement of in axon regeneration

To confirm that the K48 and K63 ubiquitination pathways, autophagy and casein kinases are involved in axon regeneration we tested inhibitors on regeneration of axotomized adult rat DRG axons and human iPSC-derived sensory neurons in vitro. All the inhibitors reduced axon regeneration of rat sensory axons, particularly the autophagy inhibitor. However only the autophagy inhibitor affected continuing growth of uncut axons, confirming that regeneration of a new growth cone requires specific mechanisms that are not necessary for continuing growth (Bradke et al., 2012) . Regeneration of human iPSC-derived sensory neurons showed a similar profile of inhibition, except that no inhibition was seen with the K63 ubiquitination inhibitor.

### Control of the PNS and CNS regeneration programmes

Elements of the PNS RAG programme may be initiated by signalling from damage responses, focal adhesion (which signal via the PI3K/Akt, Ras/Erk, Jnk, rho/rac pathways) and via mechano- transduction through Piezo/Ca^++^, and Hippo/Yap/Taz (Humphries et al., 2019; Hynes, 2002). The current study shows that signalling from activated integrins alone is sufficient to upregulate many RAG transcripts. For control of the CNS regeneration programme we identified identifed Mef2a, Runx3, E2f4, Tfeb, Yy1 as potential regulators of autophagy, trafficking, ubiqutination and ER. Runx3 is involved in sensory axon guidance and connectivity, Mef2a acts via p38 which has many functions in axon growth, Yy1 has a role in hippocampal axon growth and control of autophagy, Tfeb is involved in autophagy control and E2F4 has a role in spinal axon regeneration and autophagy (Chen et al., 2006; de Angelis et al., 2005; Du et al., 2019; Klar et al., 2015; Qiang et al., 2016).

### Alpha9 integrin-kindlin-1 driven axon regeneration does not recapitulate embryonic development

The issue of whether integrin-kindlin-driven regeneration represents a return to an embryonic pattern of expression is of obvious interest. In general, repair does not accurately recapitulate development (Harty et al., 2003; Nacu and Tanaka, 2011; Seifert et al., 2019), although basic mechanisms of cell movement are shared. In a previous study, embryonic recapitulation was marginally significant in peripheral nerve regeneration (Renthal et al., 2020). But, a recent study of cortical neurons whose axons elongated in embryonic grafts in the spinal cord showed significant embryonic recapitulation (Poplawski et al., 2020). In this study, RRHO comparisons between the α9k1 regeneration group and a previous profiling study of sensory neuron expression at two embryonic ages at E12.5 and E17.5 (Tedeschi et al., 2016) showed no correlation.

Overall, the study suggests that axon regeneration in the CNS comprises two steps. In the first priming step, various transcripts, including those identified as RAGs, are upregulated by signalling from α9k1. As a second step there is a set of transcripts involved in enabling axon growth through the CNS environment, which we found to be only upregulated in the α9k1 crush/CNS regeneration group. These are involved in transport/trafficking, ubiquitination, autophagy, endoplasmic reticulum, endomembrane organization, cytoskeletal binding, and probably enable the transport and trafficking of regeneration-associated molecules and organelles, interaction with the CNS environment, and signalling.

## METHODS

### Animal surgeries

All animal surgeries were conducted in accordance with the United Kingdom Animals (Scientific Procedures) Act 1986. In this study, adult 2-month-old male Lewis rats were used. DRG virus injection was performed according to the protocol described (Cheah et al., 2017). The three viruses used in this study were AAV5-CMV-fGFP, AAV5-CAG-α9-V5 and AVV5-CMV-kindlin1-GFP, and they were obtained from the previous integrin study (Cheah et al., 2016). Briefly, one microliter of the virus at a working titer of 2 x 10^12^ GC/ml was injected into the left C5-C8 DRGs using a custom-made 33 gauge needle syringe (Hamilton) with an infusion syringe pump (World Precision Instruments) at 0.1 μl/min. For the groups of animals with crush injury, the left C5-C8 dorsal roots were crushed using a pair of Bonn forceps (Fine Science Tools) for 3 x 10 s for each root. The animals were kept for 8 weeks for recovery, and were culled by exposure to a rising concentration of CO2. The virus-injected left C5- C8 DRGs were harvested for FACS. The spinal cords were fixed with 4% PFA for *post hoc* immunohistochemical analysis.

### Fluorescence-activated cell sorting (FACS)

Harvested DRGs were dissociated by incubating with 0.2% collagenase (Sigma) and 0.1% trypsin (Sigma), followed by trituration and centrifugation (Cheah et al., 2016). After dissociation, the cells were kept in ice-cold FACS buffer containing 1% BSA (Sigma), 2 mM EDTA (Sigma), 10mM NaN3 (Sigma), 15mM HEPES (Gibco) in PBS at pH 7.4, and immediately proceeded to FACS. The FACS Aria III Flow Cytometer (BD Biosciences) was used for cell sorting. Dissociated rat nestin-GFP left C5-C8 DRG cells were used as positive control to adjust forward and side scatters for doublet discrimination and debris exclusion. To obtain a pure population of GFP-positive cells, 488 nm emission with 530/30 BP filter was used, coupled with 633 emission and 660/20 BP filter as negative control. GFP-sorted cells were kept in ice-cold RNA*later* (Ambion) and then moved to -80°C for storage in preparation for RNA-seq library.

### RNA-seq library preparation

RNA from FACS-sorted cells ranging from 3,000 – 10,000 were isolated with the NucleoSpin RNA XS kit (Clontech) with on column DNase digestion according to the manufacturer’s protocol. The SMART-Seq v4 Ultra Low Input RNA Kit (Clontech) was used for library preparation. The cDNA was fragmented to 300 base pairs (bp) using the Covaris M220 (Covaris), and then the manufacturer’s instructions were followed for end repair, adaptor ligation, and library amplification. The libraries were quantified by the Qubit dsDNA HS Assay Kit (Molecular Probes); library size distribution and molar concentration of cDNA molecules in each library were determined by the Agilent High Sensitivity DNA Assay on an Agilent 2200 TapeStation system. Libraries were multiplexed into a single pool and sequenced using a HiSeq2500 instrument (Illumina, San Diego, CA) to generate 69 bp pair-end reads. An average of ∼60 million reads were obtained for each sample.

### RNA-seq read alignment and processing

RNA-seq data were mapped to the reference genome (mm10 / GRCm38) using STAR (Dobin et al., 2013). Aligned reads were sorted, and alignments mapped to different chromosomes were removed from the BAM file using SAMtools (http://www.htslib.org/). Alignment and duplication metrics were collected using the PICARD tools functions CollectRnaSeqMetrics and MarkDuplicates respectively (http://broadinstitute.github.io/picard/). Total counts of read fragments aligned to candidate gene regions were quantified by Salmon (Patro et al., 2017). Genes with less than 10 read counts in over 80% of the samples were removed. Gene counts were log2 – transformed with a small pseudocount. Potential bias from gene length and sequencing depth was normalized using *cqn* R package (Hansen et al., 2012).

### Differential gene expression

Principle component analysis (PCA) of the normalized expression data (first five PCs) was correlated with potential technical covariates, including sex, batch, aligning and sequencing bias calculated from STAR and Picard respectively. Differential gene expression was performed using limma (Ritchie et al., 2015) on normalized gene counts, including batch the first three PCs of aligning and sequencing bias as covariates: ∼ Genotype*Condition + Batch + AlignSeq.PC1 + AlignSeq.PC2 + AlignSeq.PC3, in which Genotype indicates mice expressing alpha9 integrin / kindlin-1 or GFP as control, and Condition indicates mice with or without dorsal root crush. Differentially expressed genes (DEGs) were determined at FDR p value < 0.1. *K*-means clustering (n = 4) were performed on genes that are up-regulated by overexpressing alpha9 integrin / kindlin-1 with or without crush, which leads to gene clusters that are uniquely or commonly up-regulated in either condition.

### Rank-Rank Hypergeometric Overlap (RRHO)

We performed a rank-rank hypergeometric test as previously described (Plaisier et al., 2010). Genes were ranked according to their logFC before running the rank–rank hypergeometric overlap test to evaluate overlap between two datasets.

### Gene Ontology Analysis

GO term enrichment analysis was performed using the gProfileR package (Reimand et al., 2016), using expressed genes in each of the normalized dataset as background. Genes enriched in the top GO terms of interest ranked by FDR *P*-value were manually checked on Uniprot and PubMed for relevance. The very general terms were excluded, and terms with low P values and potential relevance selected. GO terms with similar contents were gathered together as shown on the y-axis of the individual heatmaps in **Fig. 3A-C, S3, S4** and in the tables in **Table S1, S2** together with genes contained in these categories.

### Weighted gene co-expression network analysis

Sequencing and aligning covariates were regressed out from normalized expression data using a linear model. Co-expression network was constructed using the WGCNA package (Langfelder and Horvath, 2008). Briefly, pair-wise Pearson correlations between each gene pair were calculated and transformed to a signed adjacency matrix using a power of 10, as it was the smallest threshold that resulted in a scale-free R^2^ fit of 0.8. The adjacency matrix was used to construct a topological overlap dissimilarity matrix, from which hierarchical clustering of genes as modules were determined by a dynamic tree-cutting algorithm.

### WGCNA module annotation

To classify up- or down-regulated modules, the module eigengene, defined as the first principle component of a module that explains the maximum possible variability of that module, was related to genotype (alpha9 integrin / kindlin-1 vs GFP) using a linear model. Modules were considered to be significantly associated with the phenotype when Bonferroni corrected p values are less than 0.05. To annotate modules at a general level, we applied gene ontology (GO) enrichment analyses on each module. As a further step towards functional annotation, we performed hypergeometric analysis to examine each module’s association with the regeneration-associated gene (RAGs) module known to be activated by peripheral injury (Chandran et al., 2016), or transport-associated gene module. Modules were considered to be significantly associated with each of the gene program when Bonferroni corrected p values are less than 0.05. The Pearson correlations between each gene and each module eigengene as a gene’s module membership were also calculated (**Supplemental Table 4**), and hub genes were defined as being those with highest correlations (kME > 0.85). The co-expression networks of these most central genes representing key biological pathways up-regulated in the regenerating DRG neurons were plotted using *igraph* R package (https://igraph.org/r/). Correlations of ME between DEG clusters and WGCNA modules were calculated using *bicor()* in the WGCNA package and bi-weighted mid-correlation (R) values were plotted in a multidimensional scaling plot.

### Transcription factor enrichment analysis

*Enrichr* (Chen et la., 2013; Kuleshov et la., 2016) was used to predict the upstream transcription factors of each WGCNA modules using position weight matrices (PWMs) from TRANSFAC and JASPAR scanning the promoters of genes in the region between −2000 and +500 of a gene’s transcription start site. This workflow also integrates existing ChIP-seq data from ENCODE (Rosenbloom et al., 2012) and ChEA (Lachmann et la., 2010). The TF-target network containing predicted TFs and module genes predicted to be bound by each TF was plotted using *igraph*.

### Ubiquitination network analysis

UbiNet (Nguyen et al., 2016) was utilized to search for ubiquitination networks and hubs amongst mRNAs expressed in the Magenta/Black/Turquoise modules. The upregulated genes were submitted to the UbiNet Network Analysis platform to identify the potential E3 ligase-substrate interactions. The most interacted E3 ligases or hubs were selected, with their cellular functions cross-checked manually on UniProt and PubMed.

### Immunohistochemical analysis

Sections of PFA-fixed spinal cords were cut at 14 μm on a cryostat and blocked in 0.4% Triton X-100 (Sigma) and 10% normal goat serum (Invitrogen). The tissues were then incubated with primary antibodies overnight at 4°C and secondary antibodies for 2 hr at room temperature. Primary antibodies used were rabbit anti-GFP (1:500; Invitrogen A-11122) and mouse anti-V5 (1:250; Invitrogen R963-25); secondary antibodies were Alexa Fluor 488 and 568 (1:500; Invitrogen).

### DRG explant neurite regeneration assay

Cervical DRGs from 2-month-old male Wistar rats were isolated with connective tissues removed. Each DRG was trimmed into 2-4 pieces and plated on μ- Slide angiogenesis chamber slides (Ibidi) coated with poly-D-lysine (20 μg/ml; Sigma) and laminin (10 μg/ml; Sigma). The culture media were refreshed daily to promote neurite outgrowth. On Day 5, inhibitors were added to the media and incubated for 1 hr before axotomy. The inhibitors used were: NSC697923 (0.3 µM, Tocris Bioscience), MLN4924 (5 µM, Tocris Bioscience), MG132 (10 µM, Abcam), 3-Methyladenine (2.5 mM, Abcam) and D4476 (10 µM, Tocris Bioscience). To perform axotomy, a glass pulled pipette was used to leave a clear demarcation on the slides. The tissues were returned to the incubator for 2 hr after axotomy. Images were taken immediately after axotomy and at 2 hr using the Axio Observer D1 Inverted Phase Contrast Microscope (Zeiss). Analysis was performed using ImageJ by quantifying the number of regenerating neurites.

### Human iPSC-derived sensory neuron regeneration assay

Human iPSC-derived sensory neurons were generated in a collaboration with Ilyas Singec’s group at the National Center for the Advances of Translational Sciences (NCATS) at the NIH (Patent Publication Number: WO / 2020/219811) (Singec et al., 2020). In short, human iPSCs were cultured until 70-80% confluent, treated with a combination of small molecule inhibitors and grown into neurospheres for 14 days. Neurospheres were then dissociated and replated as individual spot cultures on 24-well plates. Each spot contained 60,000 cells and was further matured for 21 days until long axons protrude out of the spot. On day 35 DIV, cells were treated with inhibitors 2 hours before laser axotomy was performed with a 300mW Stiletto infrared laser (Hamilton Thorne) combined with a 20x objective as described previously (Ho et al., 2021). The laser module has a high-speed micro controller and an automated motorized stage. Axons were cut around 400-600 µm away from the cell bodies. Spots were imaged before and every 2 hours for up to 24 hours after cut using the live-cell imaging system – IncuCyte S3 (Satorius). Images were extracted from the InCucyte software and areas of interested were cropped at 4 inc x 6 inc including the injury site. Images were then converted into 8-bit images in Fiji ImageJ and Sholl analysis was performed at every micron for 250 um away from the injury site at different time points. % regeneration was measured by the total number of processes at time x after cut / total number of processes before cut. The inhibitors used were: NSC697923 (0.3 µM, Tocris Bioscience), MLN4924 (5 µM, Tocris Bioscience), MG132 (10 µM, Abcam), 3-Methyladenine (0.5 mM, Abcam) and D4476 (10 µM, Tocris Bioscience).

## AUTHOR CONTRIBUTIONS

MC, YC, VP, VS, CJW, DHG and JWF designed experiments; MC, YC, VP, AC, CJW, DHG and JWF analyzed data; MC, YC, VP and AC performed experiments; MC, YC, DHG and JWF wrote the paper; CJW, DHG and JWF secured funding.

## ACCESSION NUMBERS

The accession numbers for the data generated in this paper are GSE188775 and GSE188776.

## ACKNOWLEDGEMENT

This work was supported by grants from Medical Research Council (MR/R004544/1 and MR/R004463/1), NWO (013-16-002), Fight for Sight (5065/5066, 5119/5120 and 5123/5124), Wings for Life (WFL GB-04/19), Czech Ministry of Education (CZ.02.1.01/0.0./0.0/15_003/0000419), ERA-NET NEURON AxonRepair, International Foundation for Research in Paraplegia P172, National Institutes of Health R35NS105076, and Dr Miriam and Sheldon G Adelson Medical Research Foundation.

---

**Figure S1.**
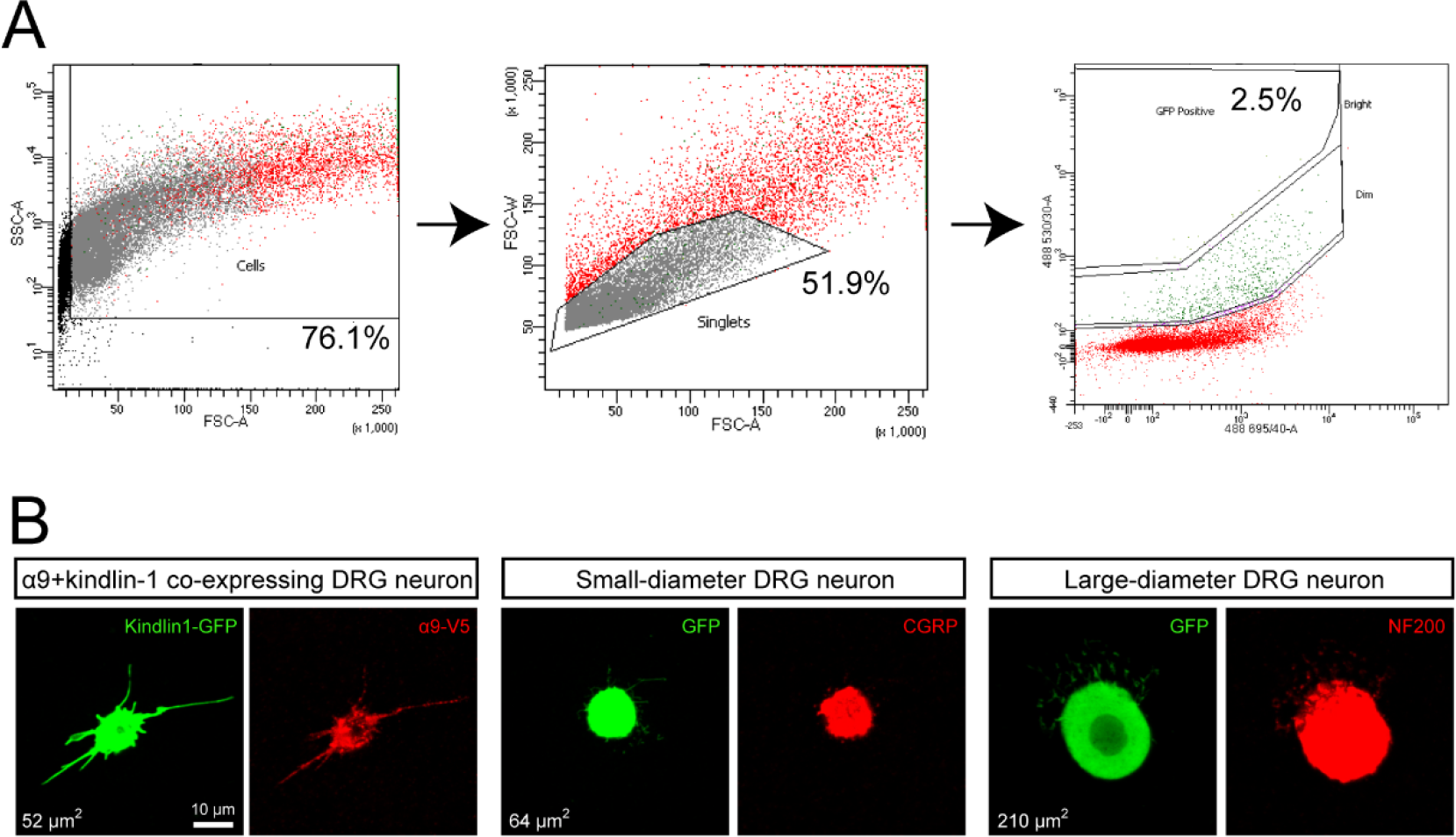
FACS sorting of GFP-expressing DRG neurons. **A** shows multi-gated approach of sorting dissociated GFP-expressing DRG neurons using the BD Aria III Flow Cytometer. Dissociated rat nestin-GFP left C5-C8 DRG cells were used as positive control to adjust forward and side scatters for doublet discrimination and debris exclusion. A pure population of single dissociated GFP-expressing DRG neurons, approximately 2.5% of the total cell population, was obtained. **B** shows a small sample of sorted DRG neurons were cultured and analysed for α9 integrin and kindlin-1 expression, and co-stained with CGRP and or NF200 for verification.

**Figure S2.**
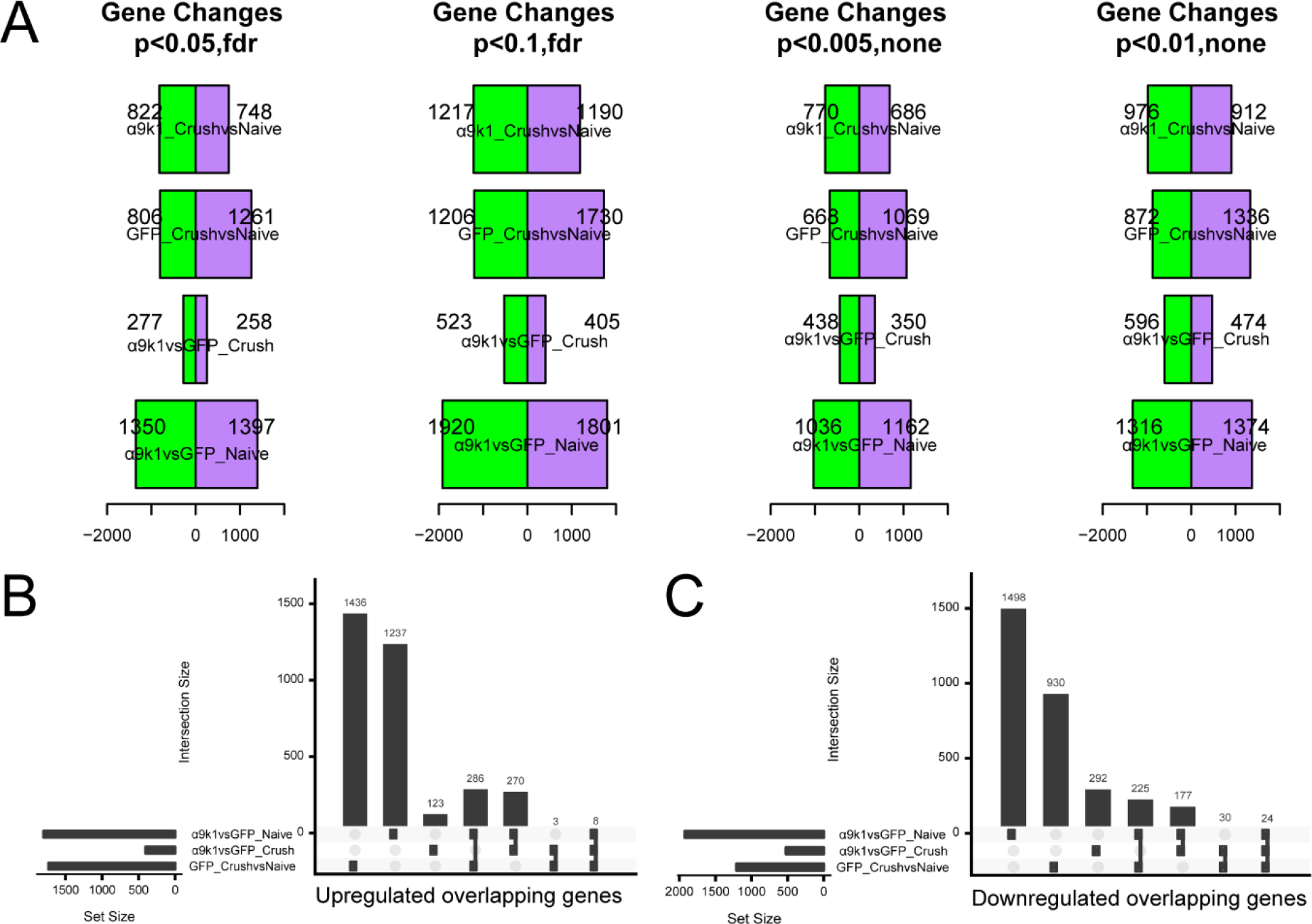
Differential expression between experimental and control groups. **A** shows the number of differentially expressed genes in different experimental and control groups: α9k1-crush vs α9k1-naive, GFP-crush vs GFP-naive, α9k1-crush vs GFP-crush, and α9k1-naive vs GFP- naïve. **B-C** show quantitation of upregulated (**B**) and downregulated (**C**) overlapping genes from **A**.

**Figure S3.**
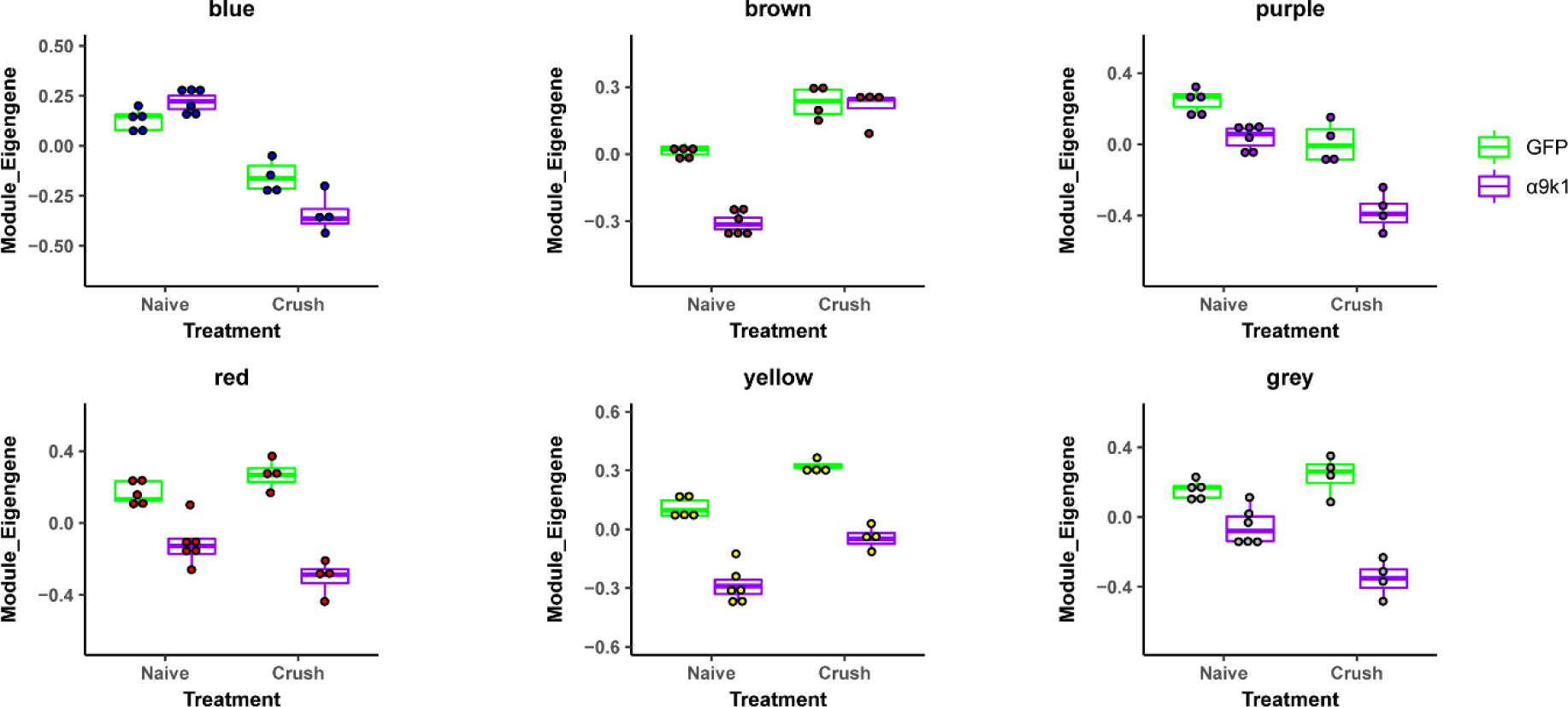
Irrelevant gene modules to our analysis due to the lack of correlation to any specific treatment groups of interest.

**Figure S4.**
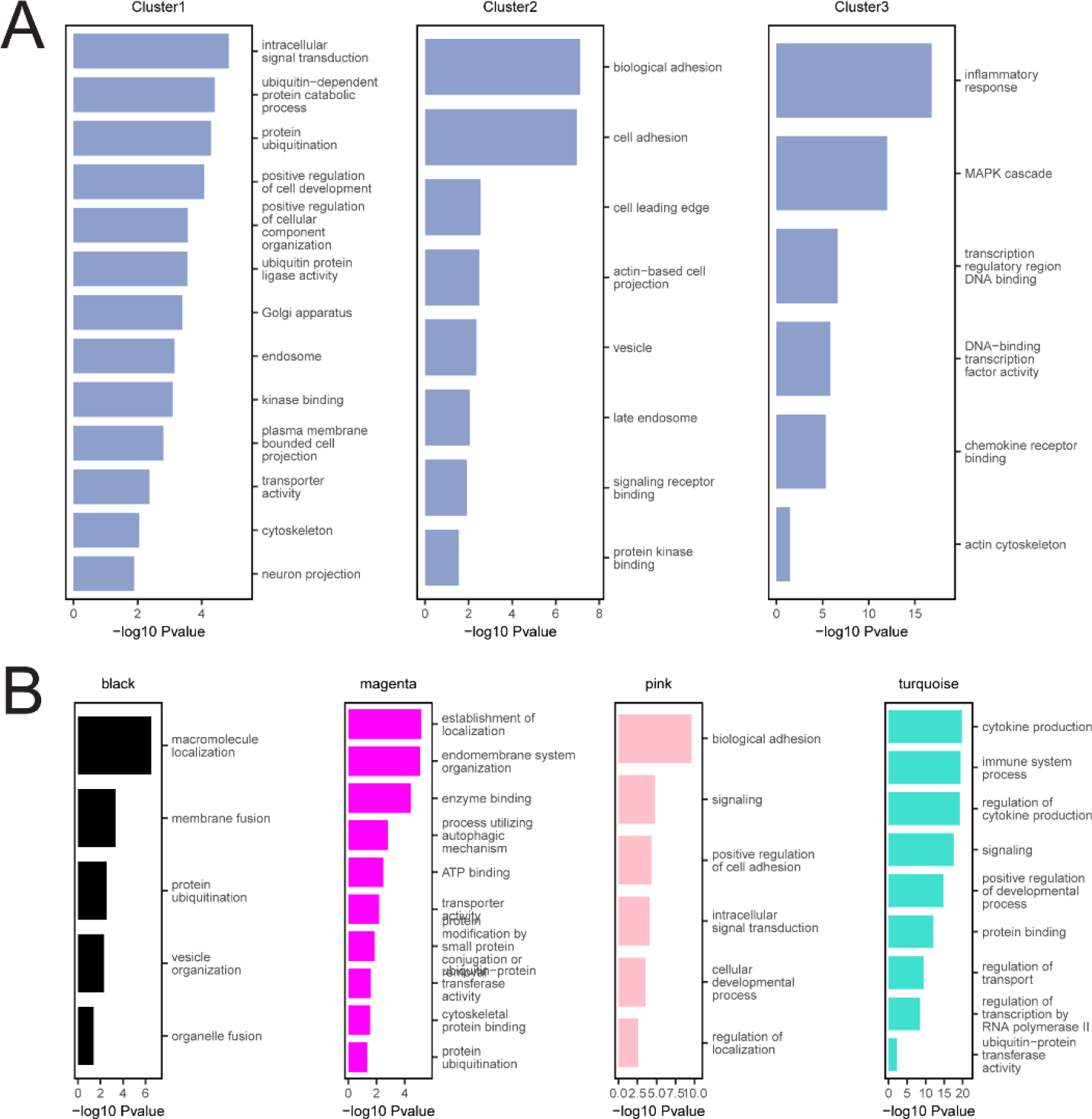
Top GO terms for Clusters 1-3, Black, Magenta, Pink and Turquoise modules. **A-B** show top GO terms of interest ranked by *P*-value for Clusters 1-3 (**A**) and the 4 colored modules (**B**) using GO terms with an enrichment FDR P < 0.05 (Methods). Genes enriched in the top GO terms were manually checked on UniProt and PubMed for relevance.

**Figure S5.**
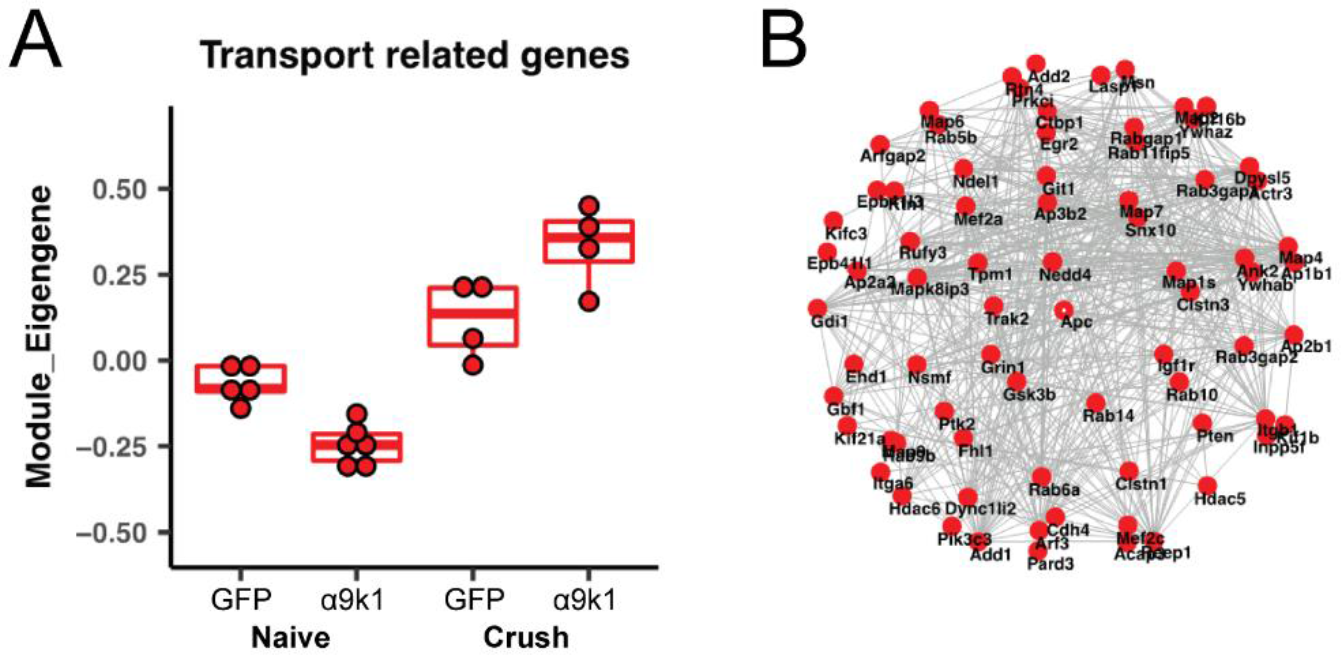
Transport and trafficking-related molecules in Black, Magenta, Turquoise and Pink modules. **A** shows the α9k1-crush group upregulated the highest degree of transport-related molecules in the four modules which play a role in axon regeneration. **B** shows the top 100 most connected or hub genes related to transport. These genes are also enriched in the RAG_Chandran and Transport_Koseki datasets in Figure 6A.

**Figure S6.**
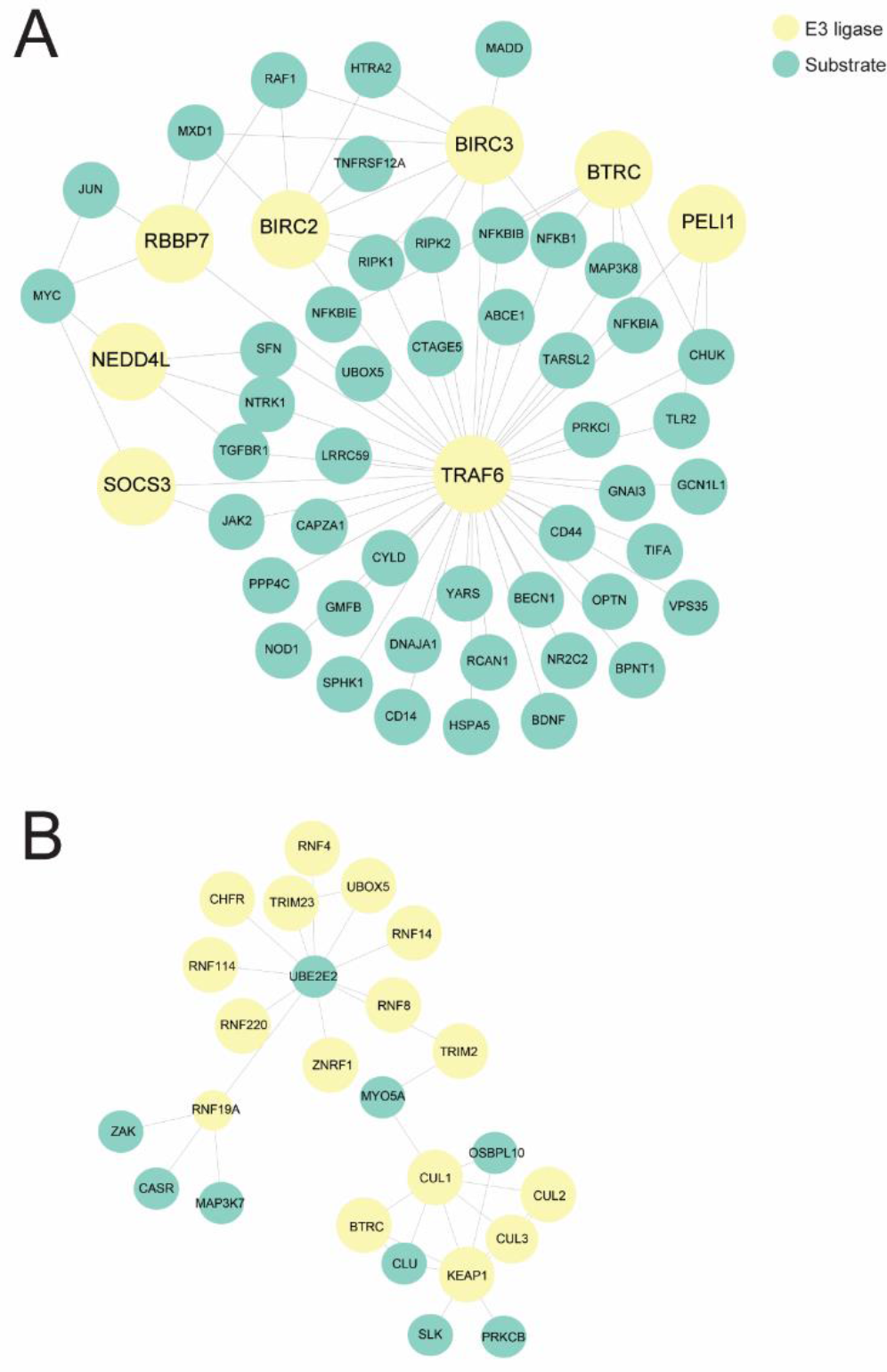
Additional ubiquitination network analysis related to sensory axon regeneration using UbiNet. **A.** A network centres around the hub Traf6, an E3 ligase implicated in promoting tumorigenesis and invasion through activation of AKT signalling and in control of the NF-kappa-B family. **B.** A network with mostly Ube2e2 and Keap1 as hubs which is associated with oxidative stress and neuroprotection.

**Table S1.**
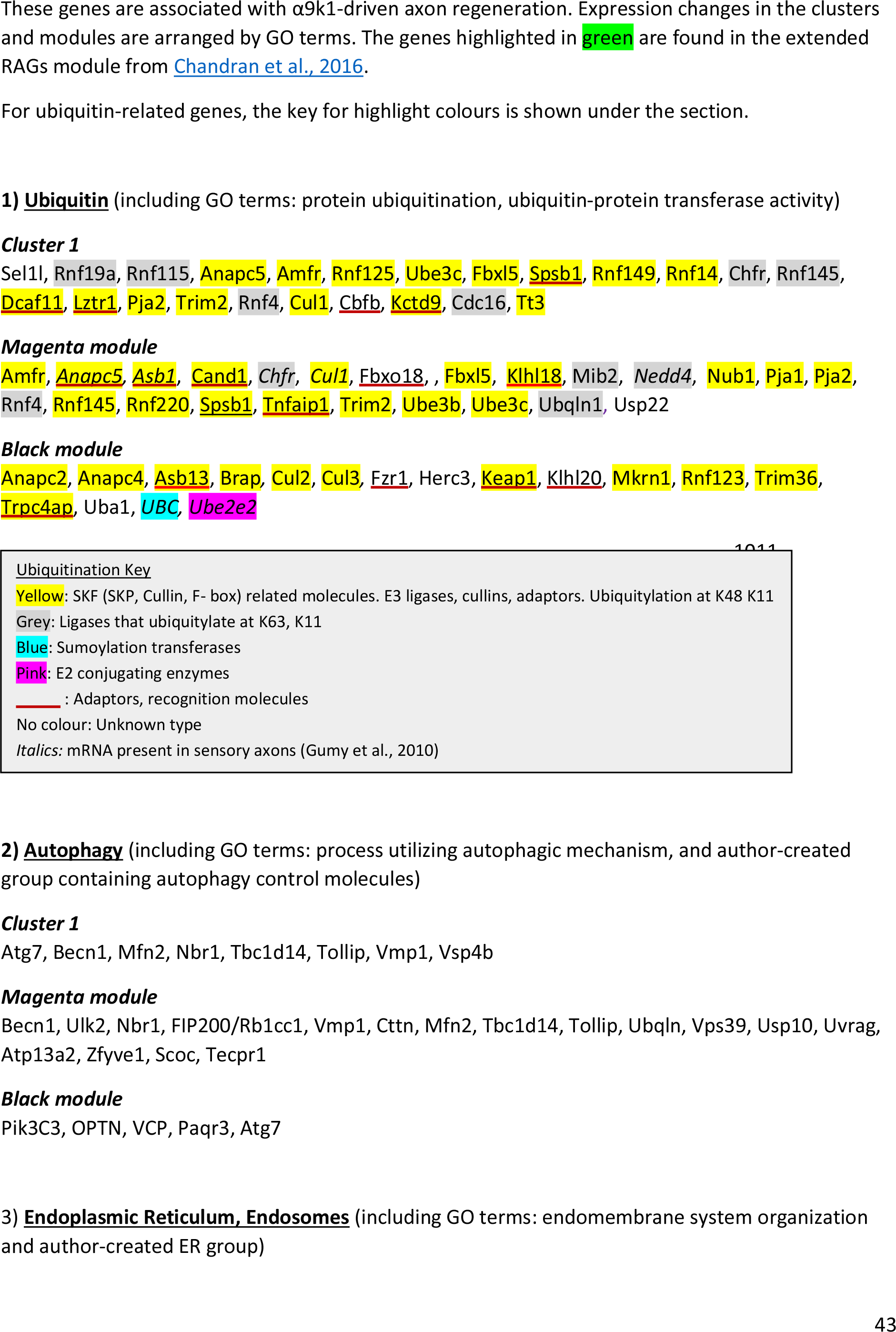
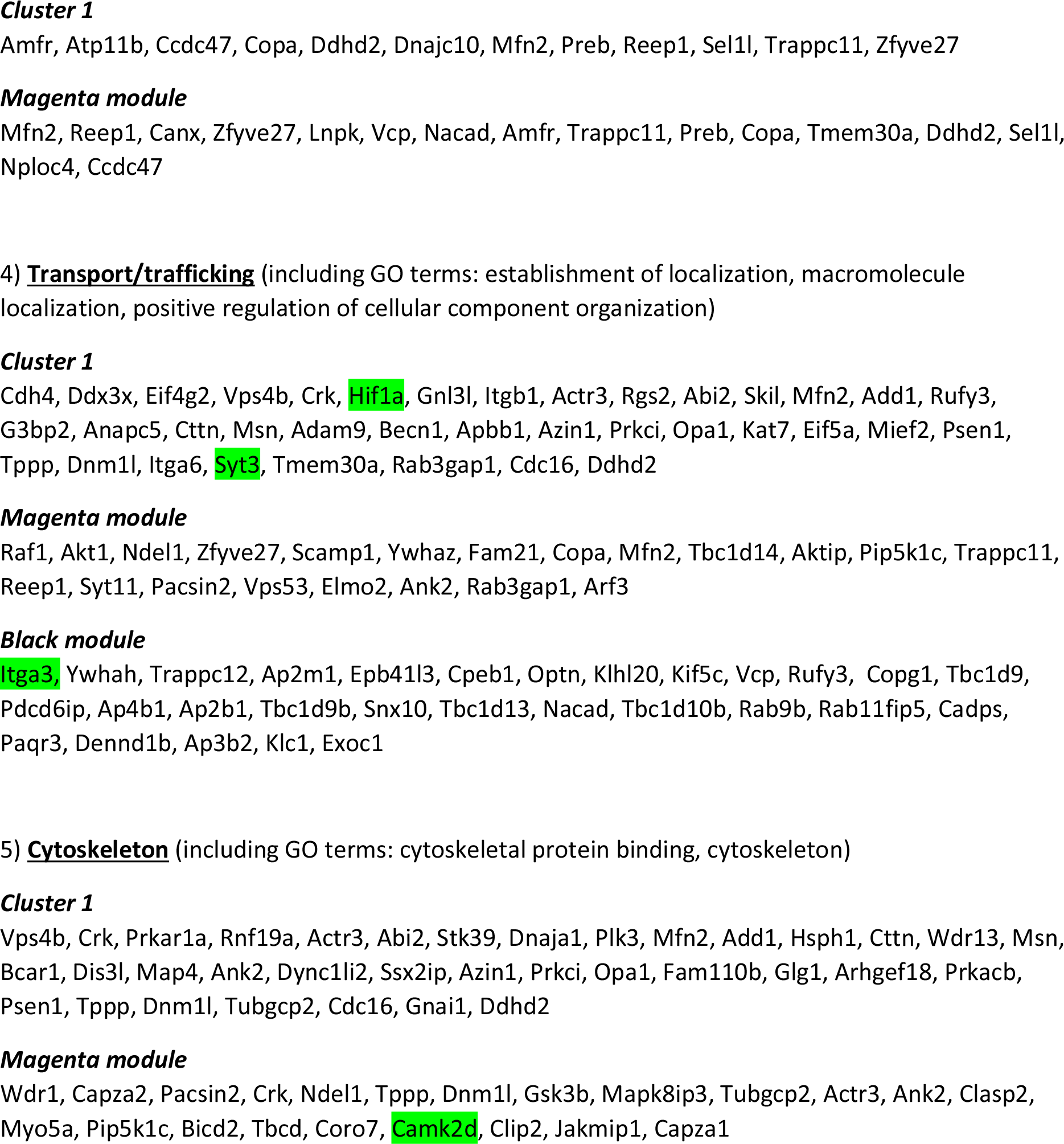
Genes upregulated in Cluster 1, Magenta and Black modules for the α9k1-crush (regeneration) group

**Table S2.**
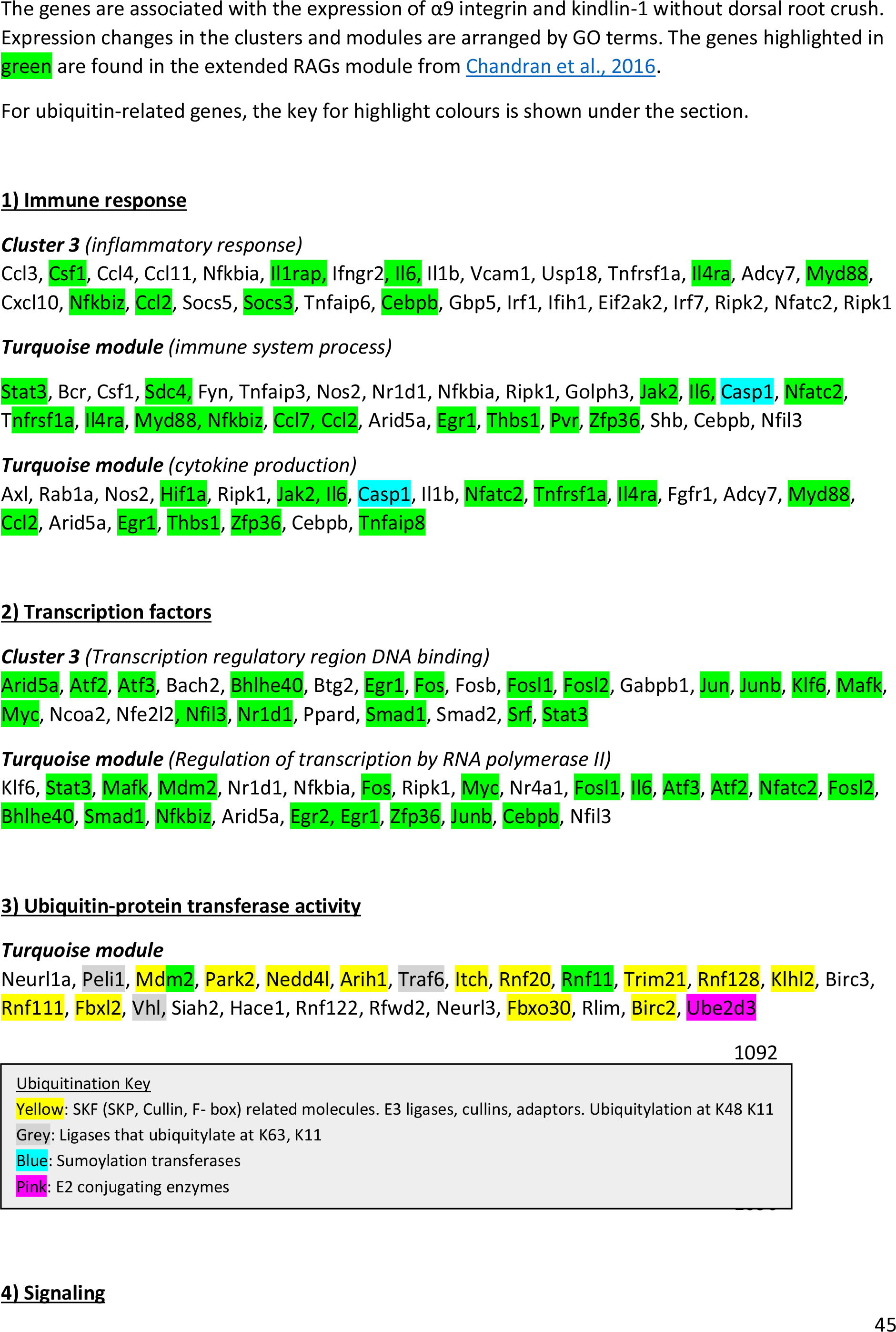
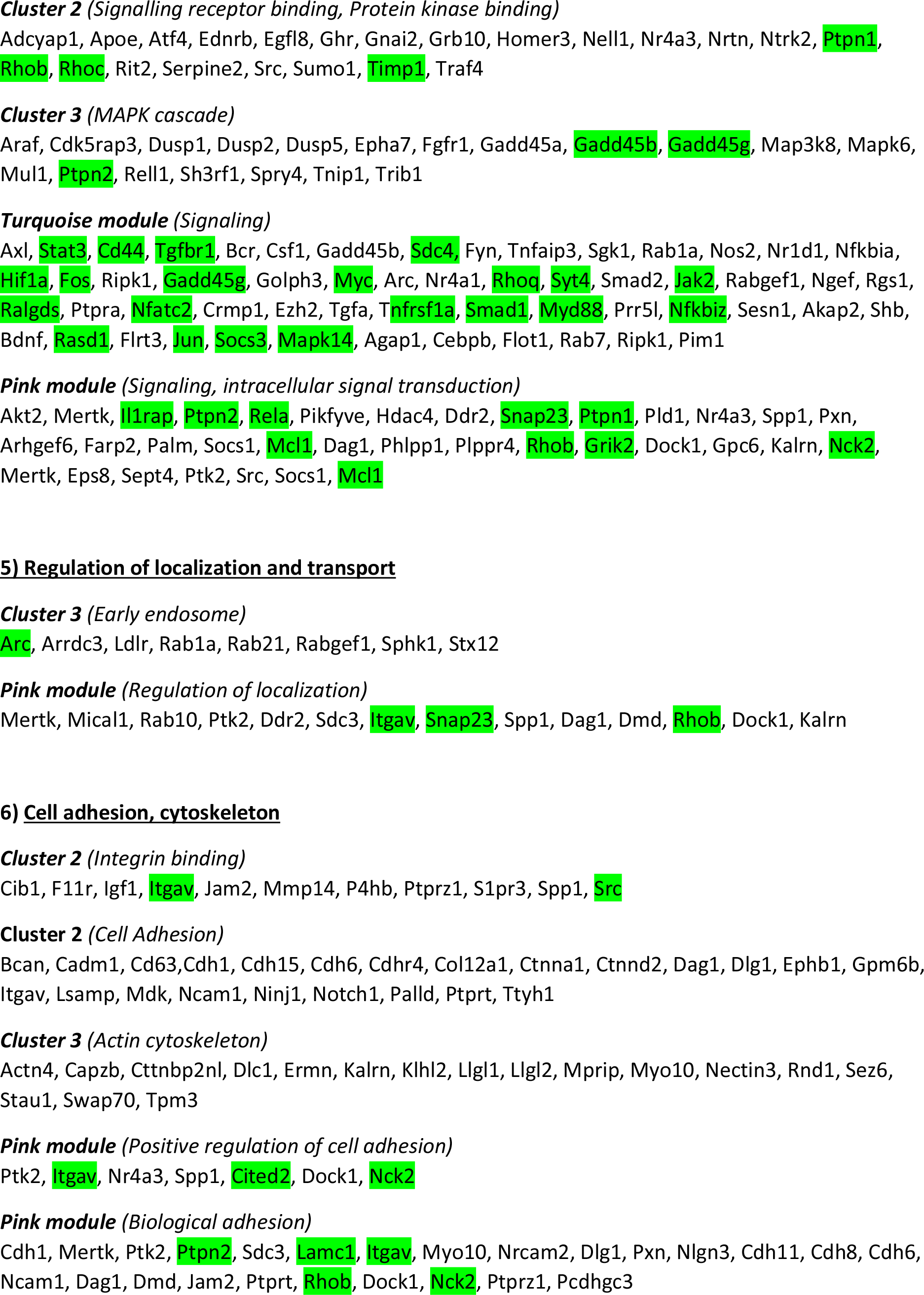
Genes upregulated in Clusters 2 and 3, Pink and Turquoise modules for the α9k1-naïve group

